# The deubiquitinase Rpn11 functions as an allosteric ubiquitin sensor to promote substrate engagement by the 26S proteasome

**DOI:** 10.1101/2024.10.24.620116

**Authors:** Zaw Min Htet, Ken C. Dong, Andreas Martin

**Affiliations:** Department of Molecular and Cell Biology, University of California at Berkeley, Berkeley, CA 94720, USA; California Institute for Quantitative Biosciences, University of California at Berkeley, Berkeley, CA 94720, USA; Howard Hughes Medical Institute, University of California at Berkeley, Berkeley, CA 94720, USA

## Abstract

The 26S proteasome is the major compartmental protease in eukaryotic cells, responsible for the ATP-dependent turnover of obsolete, damaged, or misfolded proteins that are delivered for degradation through attached ubiquitin modifications. In addition to targeting substrates to the proteasome, ubiquitin was recently shown to promote degradation initiation by directly modulating the conformational switching of the proteasome, yet the underlying mechanisms are unknown. Here, we used biochemical, mutational, and single-molecule FRET-based approaches to show that the proteasomal deubiquitinase Rpn11 functions as an allosteric sensor and facilitates the early steps of degradation. After substrate recruitment to the proteasome, ubiquitin binding to Rpn11 interferes with conformation-specific interactions of the ubiquitin-receptor subunit Rpn10, thereby stabilizing the engagement-competent state of the proteasome and expediting substrate insertion into the ATPase motor for mechanical translocation, unfolding, and Rpn11-mediated deubiquitination. These findings explain how modifications with poly-ubiquitin chains or multiple mono-ubiquitins allosterically promote substrate degradation and allow up to four-fold faster turnover by the proteasome.

## Introduction

The 26S proteasome plays indispensable roles for eukaryotic cell function and viability by carrying out the targeted degradation of misfolded, damaged, or aggregated proteins for general homeostasis and quality control, as well as the turnover of numerous regulatory proteins involved in vital cellular processes ^1,2^. The proteasome must therefore be able to selectively degrade appropriate substrates in a crowded environment and in response to dynamic cellular needs, yet possess high promiscuity to process diverse polypeptide structures and sequences. This is achieved by a bipartite degradation signal combined with the intricate architecture of the proteasome and its switching between conformationally distinct states. For the majority of proteasomal substrates, the first component of the bipartite degradation signal constitutes a ubiquitin modification that is covalently conjugated to one or several lysine residues and targets the substrate to a proteasomal ubiquitin receptor. *In vivo* studies have identified homotypic K48- and K11-linked polyubiquitin chains as the primary proteasomal targeting signals ^3,4^ that interact with one or several of the three main ubiquitin receptors, Rpn1, Rpn10 and Rpn13 ^5,6^ ^7,8^. In addition, other proteasome subunits such as Rpt5 and Sem1 were shown to bind ubiquitin without being *bona fide* ubiquitin receptors required for proteasomal substrate degradation ^9^ ^10^. The second component of the degradation signal is an unstructured initiation region of sufficient length and complexity to be inserted into the proteasome’s central channel for stable engagement by the six ATPase subunits of the AAA+ (ATPases Associated with diverse cellular Activities) motor^11 4,12–18^.

The proteolytic active sites of the 26S proteasome are sequestered inside a barrel-shaped 20S core particle, which is capped on one or both ends by a 19S regulatory particle that can be further subdivided into the lid and base subcomplexes ^11^. The lid contains several structural subunits and the essential deubiquitinase Rpn11 ^19,20^, while the base includes the ubiquitin receptors, Rpn1, Rpn10, and Rpn13, and the six distinct ATPase subunits Rpt1-Rpt6 that form a heterohexameric ATPase motor ^8,11,21^. After substrate recruitment through ubiquitin binding to one or several of the receptors, the flexible initiation region of the substrate diffuses into the central channel of the AAA+ motor, which is made up from an N-terminal domain ring (N-ring) sitting on top of an ATPase-domain ring (Supplementary Fig. 1) ^12^. Conserved pore loops project from each ATPase domain of the hexameric ring into the central channel, sterically engage the substrate, and transduce ATP-hydrolysis-driven conformational changes of the ATPase ring for mechanical unfolding and translocation of the substrate polypeptide into the 20S core peptidase ^22,23^. Rpn11 is located above the N-ring and catalyzes the *en-bloc* removal of substrate-attached ubiquitins prior to their entry into the central channel ^19,24^.

Substrate insertion into the ATPase motor and engagement with the pore loops is further regulated by the conformational switching of the proteasome ^25^. Previous structural studies revealed several conformations that can be grouped into two main classes, the engagement-competent (s1) state and the processing-competent (non-s1) states, which exist in a dynamic equilibrium (Fig. 1c) ^26–35^. In the absence of substrate, the proteasome primarily adopts the engagement-competent s1 state, in which the entrance to the N-ring is well accessible for substrates, but the central channel through the N-ring and the ATPase ring is not coaxially aligned with the core particle. By contrast, in the processing-competent non-s1 states, a wider, coaxially aligned channel throughout the regulatory particle and into the 20S core facilitates processive substrate translocation, and a centrally localized Rpn11 obstructs the entrance to this channel, yet enables the efficient co-translocational deubiquitination of a substrate that was previously inserted. The proteasome therefore must adopt the engagement-competent s1 state for insertion of a substrate’s flexible initiation region, before successful engagement by the ATPase motor triggers the conformational switch to the processing-competent non-s1 states for translocation, deubiquitination, and mechanical unfolding ^12,36^. Our previous single-molecule FRET-based studies revealed that the ATP-hydrolyzing, substrate-free 26S proteasome spontaneously switches between s1 and non-s1 states, with a forward s1 - to - non-s1 transition rate that is about 4-fold lower than the backward non-s1 - to - s1 rate, leading to a predominance of the engagement-competent s1 state ^36^. Selective destabilization of the s1 state through the disruption of interactions at the lid-base interface consequently leads to major defects in substrate engagement, while later substrate-processing steps remain unaffected ^37^. Modulating the conformational switching of the proteasome and thereby substrate access to the degradation machinery may thus play important roles for substrate selectivity and prioritization.

**Fig. 1:**
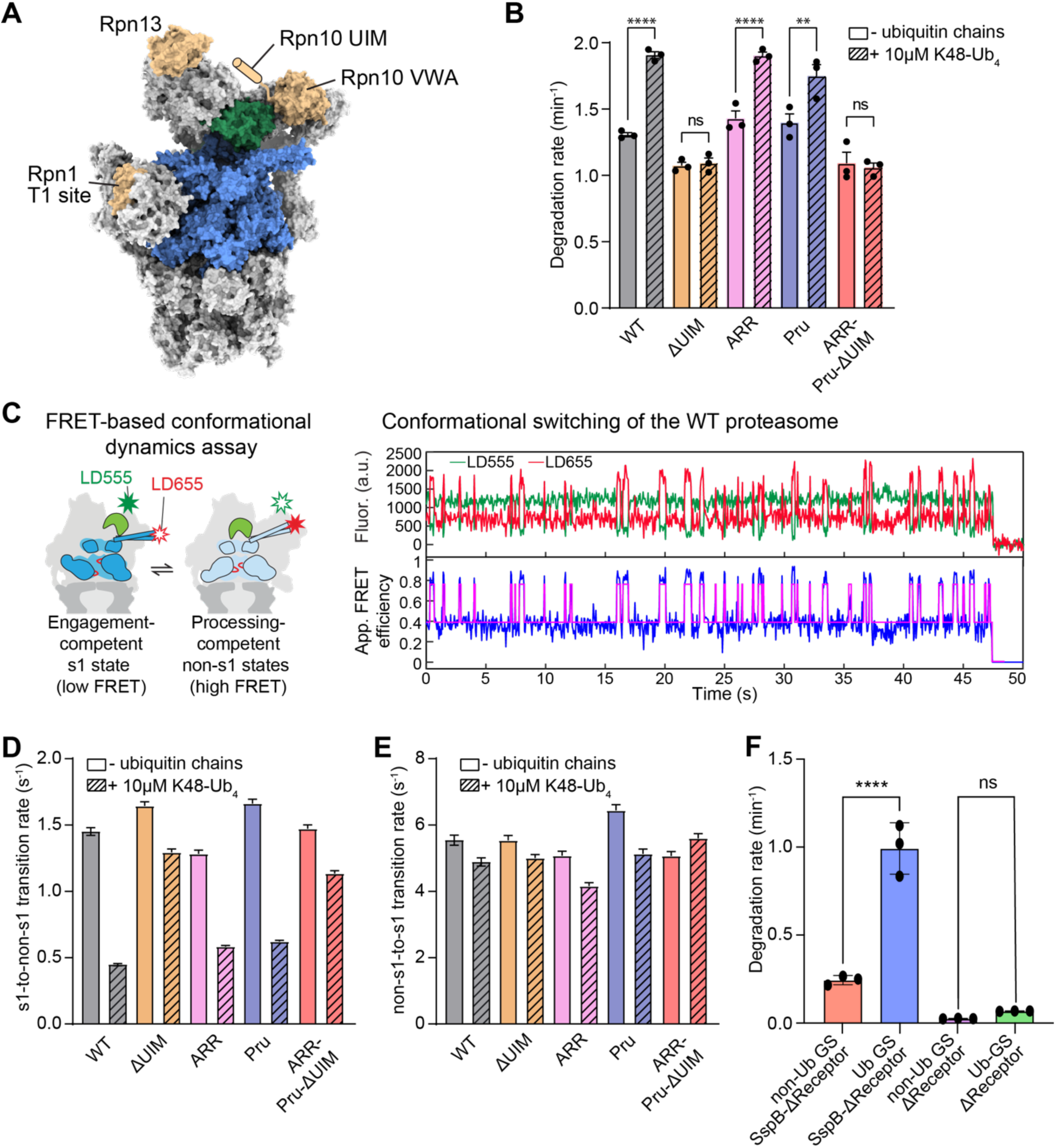
The ubiquitin interacting motif of Rpn10 mediates the allosteric effect of ubiquitin chains on the proteasome conformational dynamics and substrate degradation. A) Architecture of the 26S proteasome (PDB ID: 4CR2) with the AAA+ motor shown in blue, Rpn11 in green, and the ubiquitin receptors, Rpn1, Rpn10, and Rpn13 in tan. Rpn10’s flexible ubiquitin interacting motif (UIM) is not resolved in cryo-EM structures and depicted as a cartoon in its approximate location. B) Unanchored K48-Ub_4_ ubiquitin-chain dependent stimulation of SspB-delivered substrates degradation by receptor-deficient 26S proteasomes, either lacking the UIM of Rpn10 (ΔUIM), carrying mutations in the T1 site of Rpn1 (ARR), mutations in the Pru domain of Rpn13 (Pru) or with a combination of all these mutations (ARR-Pru-ΔUIM). Shown are the averages of three technical replicates with error bars representing the standard errors of the mean. Statistical significance was calculated using an ordinary one-way ANOVA test. ns non-significant with p>0.9999, ** p=0.0035, **** p<0.0001. For proteasomes with two mutant receptors see Supplementary Fig. 3A. C) Left: Schematic of the single-molecule FRET-based conformational dynamics assay, with a donor dye attached to Rpn9 and an acceptor dye on Rpt5, reading out the conformational switching between the s1 state (low FRET efficiency) and non-s1 states (high FRET efficiency). Right: Example traces for the donor and acceptor fluorescence (top) and the calculated apparent FRET efficiency (bottom) during the spontaneous conformational switching of an immobilized 26S proteasome in ATP. The magenta line represents the two-state fit using hidden Markov modeling. D) Effects of unanchored K48-Ub_4_ on the s1-to-non-s1-transition rates for wild-type and receptor-deficient proteasomes. Shown are the transition rates calculated by fitting the s1-state dwell time distribution observed in at least 200 FRET-efficiency traces from two technical replicates of the proteasome conformational dynamics assay, with error bars representing the standard errors of the fit. For proteasomes with two mutant receptors see Supplementary Fig. 3B. E) Effects of unanchored K48-Ub_4_ on the non-s1-to-s1 transition rates for wild-type and receptor-deficient proteasomes. Shown are the transition rates calculated by fitting the non-s1-state dwell time distribution observed in at least 200 FRET-efficiency traces from two technical replicates of the proteasome conformational dynamics assay, with error bars representing the standard errors of the fit. For proteasomes with two mutant receptors see Supplementary Fig. 3C. F) Degradation of the slowly engaging GS substrate with glycine-serine-rich initiation region in its ubiquitinated (Ub-GS) or non-ubiquitinated form (non-Ub GS) by the triple-receptor-deficient (ARR-Pru-ΔUIM) proteasome with SspB fusion (SspB-ΔReceptor) or without SspB fusion (ΔReceptor). Shown are the average rates from three technical replicates, with error bars indicating the standard deviation. Statistical significance was calculated using an ordinary one-way ANOVA test: **** p<0.0001, ns non-significant with p=0.8928.

Ubiquitin signals have recently been discovered to affect substrate-degradation kinetics ^38,39^ and regulate the conformational dynamics of the proteasome. Cryo-EM structures of the proteasome show an effect of K48-linked ubiquitin chains on the distribution of conformational states ^35^. More importantly, our previous single-molecule studies revealed that binding of ubiquitin chains allosterically stabilizes the proteasome in the engagement-competent s1 state through a ∼ 3-fold reduction of the s1 - to - non-s1 transition rate, which promotes substrate insertion and engagement with the pore loops, and consequently facilitates degradation ^36^. Despite these previous insights, the mechanistic basis for how ubiquitin chains regulate the conformational dynamics of the proteasome remains unclear.

Here, we used single-molecule FRET measurements and mutational studies of the recombinant yeast 26S proteasome to investigate how different ubiquitin-interacting subunits of the proteasome contribute to the ubiquitin-mediated stabilization of the engagement-competent state. We found that Rpn11, prior to its role in co-degradational substrate deubiquitination, functions as a ubiquitin-dependent allosteric sensor to promote initial substrate engagement by the ATPase motor, while the neighboring Rpn10 receptor acts as an important contributor to this effect by facilitating ubiquitin recruitment to Rpn11. In addition, our results revealed the interface between Rpn11, Rpt5, and Rpn10 as the regulation site that is used by ubiquitin to attenuate the conformational switching of the proteasome. This also provides a new model to further explore how the substrate geometry, the position of ubiquitin modifications, and the mode of substrate delivery to the proteasome affect substrate selectivity and degradation kinetics.

## Results

### Rpn10’s ubiquitin-interacting motif contributes to an allosteric sensor

To determine how ubiquitin modulates the proteasome conformational switching and stimulates substrate degradation, we dissected whether one of the three main ubiquitin receptors of the proteasome, Rpn1, Rpn10 and Rpn13, may be important for this effect (Fig. 1A). We used well-characterized mutations in these receptors to abrogate their ubiquitin binding: deletion of Rpn10’s ubiquitin interacting motif (UIM) (Rpn10-ΔUIM ^23^), triple point mutations D451A, D548R, and E552R in the T1 site of Rpn1 (Rpn1-ARR ^8^), and quintuple point mutations E41K, E42K, L43A, F45A, and S93D in the Pleckstrin-like receptor for ubiquitin (Pru) domain of Rpn13 (Rpn13-Pru ^8^). We first determined the allosteric effects of unanchored K48-linked tetraubiquitin chains (K48-Ub_4_) on proteasomal degradation of our titin model substrate, which consists of a N-terminal titin-I27 domain carrying the destabilizing V15P mutation and a C-terminal cyclin-B-derived unstructured initiation region with an internal ssrA tag (see Supplementary Figure 2A ^12^). To separate the effects of ubiquitin chains on substrate degradation from their role in substrate targeting, we utilized a previously established ubiquitin-independent substrate-delivery system, in which the ssrA-binding cofactor SspB from *E. coli* is fused to the N-terminus of the Rpt2 ATPase subunit of the proteasome, allowing the recruitment of non-ubiquitinated, ssrA-tagged substrates ^36,40^ (Supplementary Fig. 2A). As shown before ^36^, K48-Ub_4_ chains accelerated the SspB-delivered substrate degradation of the wildtype proteasome by ∼1.5 fold in bulk (Fig. 1B, Supplementary Fig. 2B). Using proteasomes with mutations in one of the ubiquitin receptors (ΔUIM, ARR, or Pru), we found that deletion of the Rpn10’s UIM almost completely eliminated the stimulation by unanchored K48-Ub_4_ chains on substrate degradation, whereas mutations in Rpn1 or Rpn13 did not change the ubiquitin-mediated acceleration of substrate degradation (Fig. 1B).

To further understand how ubiquitin receptors affect the proteasome conformation, we turned to our single-molecule FRET-based conformational change assay (Figure 1C), which we previously used to show that ubiquitin-induced attenuation of the engagement-competent s1 state results in faster substrate degradation ^36^. In this assay, LD555 and LD655 fluorophores are attached to position 2 of the lid subunit Rpn9 and position 49 of the Rpt5 ATPase subunit, respectively. These fluorophore placements result in low apparent FRET efficiency (∼ 0.4) for the engagement-competent s1 state and high apparent FRET efficiency (∼ 0.8) for the processing-competent non-s1 states (Fig. 1C). In the presence of ATP, wild-type proteasome primarily exists in the low-FRET s1 state with brief excursions into high-FRET non-s1 states (Fig. 1C). Consistent with our previous study, we found that ubiquitin chains decreased the rate of s1 - to - non-s1 transitions of the wildtype proteasome by ∼ 69%, from 1.45 s^−1^ to 0.45 s^−1^, whereas the rate of the non-s1 - to - s1 transition remained largely unchanged (Fig. 1D, Supplementary Fig. 4a, Supplementary Table 1)^36^. Deleting the UIM of Rpn10 significantly reduced the effect of ubiquitin chains on the s1 - to -non-s1 transition rate, whereas mutating the ubiquitin-binding sites of Rpn1 and Rpn13 did not change the ubiquitin-mediated stabilization of the s1 state (Fig. 1D, Supplementary Table 1). Similar to the wildtype proteasome, all receptor-mutant proteasome variants showed only a minor response to unanchored K48-Ub_4_ chains in their transition rate form non-s1 to s1 states (Fig. 1E, Supplementary Fig. 4B, Supplementary Table 1). These data are thus in agreement with the trends observed in our bulk degradation measurements (Figure 1B), indicating that ubiquitin binding in particular to Rpn10’s UIM significantly contributes to holding the proteasome in a conformation amenable to substrate engagement.

Multivalent interactions of ubiquitin chains with different ubiquitin receptors were previously suggested to function as a chain-length sensor ^41^, and it is therefore possible that the bridging of ubiquitin receptors with chains of sufficient length mediates the attenuation of the proteasome conformational switching and consequently affects substrate degradation. To test this model, we generated proteasome variants with mutations in two or all three of its ubiquitin receptors (ARR-Pru, Pru-ΔUIM, ARR-ΔUIM, ARR-Pru-ΔUIM) and examined the effects of ubiquitin chains on substrate degradation and conformational switching. For proteasome variants with mutations in Rpn1 and Rpn13 (ARR-Pru), K48-Ub_4_ chains affected the kinetics of SspB-delivered substrate degradation and the transition rate from the s1 to non-s1 states similar to the wildtype proteasome (Supplementary Fig. 4, Supplementary Table 1), indicating that the UIM of Rpn10 is necessary and sufficient to mediate the effect of ubiquitin chains on substrate degradation and the proteasome conformational switching.

However, we noticed that even the receptor-deficient (AAR-Pru-ΔUIM) proteasome showed a small yet significant effect of K48-Ub_4_ on the s1 - to - non-s1 transition rate (Fig. 1D, Supplementary Table 1), suggesting that Rpn10’s UIM is important for high-affinity ubiquitin binding to the proteasome, but does not represent the allosteric ubiquitin sensor responsible for controlling the conformational switching. To examine this further, we switched to a previously described slowly engaging substrate (GS substrate) that contains glycines and serines as the last 11 residues in its C-terminal unstructured tail and better mimics physiological substrates with shorter, slippery, or overall less ideal initiation regions ^12,36,42^. This substrate shows compromised initiation by the proteasomal translocation machinery and a stronger ubiquitin-mediated acceleration of degradation in the SspB delivery system ^36^. SspB-fused AAR-Pru-ΔUIM proteasomes degraded the non-ubiquitinated GS substrate at a rate of 0.25 min^−1^, and the degradation rate increased to 1.0 min^−1^ upon attachment of long ubiquitin chains (Fig. 1F, supplementary Fig. 2C). This 4-fold acceleration is thus independent of any canonical ubiquitin receptors, and an additional allosteric sensor appears to interact with the ubiquitin chains that are brought to the proteasome through the SspB-bound ubiquitinated substrate. Importantly, adding unanchored K48-Ub_4_ chains did not significantly accelerate the degradation of the non-ubiquitinated GS substrate by SspB-fused AAR-Pru-ΔUIM proteasomes (see supplementary Figure 2D), indicating that the allosteric sensor on its own binds ubiquitin only weakly and the observed small deceleration of the s1-to-non-s1 transition rate by K48-Ub_4_ in the absence of canonical ubiquitin receptors is not sufficient to facilitate the tail insertion of the slowly-engaged GS substrate. The allosteric sensor thus appears to rely on ubiquitin recruitment through intrinsic receptors, in particular Rpn10’s UIM, or through SspB binding of a ubiquitinated substrate in our artificial delivery system. This sensor thus modulates the proteasome conformational switching and substrate engagement, while Rpn10’s UIM mediates the ubiquitin interaction.

### Contacts between Rpn10 and Rpt5 regulate proteasome conformational dynamics and substrate degradation

To identify the allosteric sensor and the interfaces involved in regulating the conformational switching of the proteasome, we consulted previous cryo-EM studies. Five structures of the human 26S proteasome in the s1-like states ^43,44^ show density for ubiquitin bound to Rpn11 (Fig. 2A). Intriguingly, this ubiquitin appears to interact with the Rpt5 coiled coil and interfere with contacts formed between the same region of Rpt5 and Rpn10’s von Willebrand factor type A (VWA) domain in the processing-competent non-s1 states (Fig. 2B, C). We therefore hypothesized that ubiquitin chains stabilize the engagement-competent s1 state of the proteasome by competing with and hence preventing these non-s1 conformation-specific contacts between Rpn10 and Rpt5. To test this, we first determined whether the Rpt10-Rpt5 contacts indeed regulate substrate degradation and the proteasome conformational switching. Although the resolutions for cryo-EM structures of the *S. cerevisia*e 26S proteasome were not sufficient to unambiguously determine exact interfaces, we were able to deduce three residues in the VWA domain of yeast Rpn10, R23, D31, and E68, that may electrostatically interact with Rpt5’s coiled coil when in non-s1 states (Fig. 2C). We predicted that charge-swap mutations of these residues would therefore destabilize the non-s1 states in the absence of substrate, shift the conformational equilibrium to the s1 state, consequently show faster substrate-degradation kinetics than the wildtype proteasome, and mimic the effects of unanchored ubiquitin chains. Indeed, proteasomes containing Rpn10^EKK^ (R23E, D31K, E68K) degraded substrates ∼ 1.6-fold faster in our SspB-delivered degradation assay (Fig. 2D) and exhibited a ∼ 40% slower transition rate from the s1 to non-s1 states (Fig. 2E), with only a minor acceleration of the non-s1 - to - s1 transition (Fig. 2F, Supplementary Table 1). These results suggest that interactions between Rpn10’s VWA domain and Rpt5’s coiled-coil regulate the proteasome conformational switching and thus degradation initiation.

**Fig. 2:**
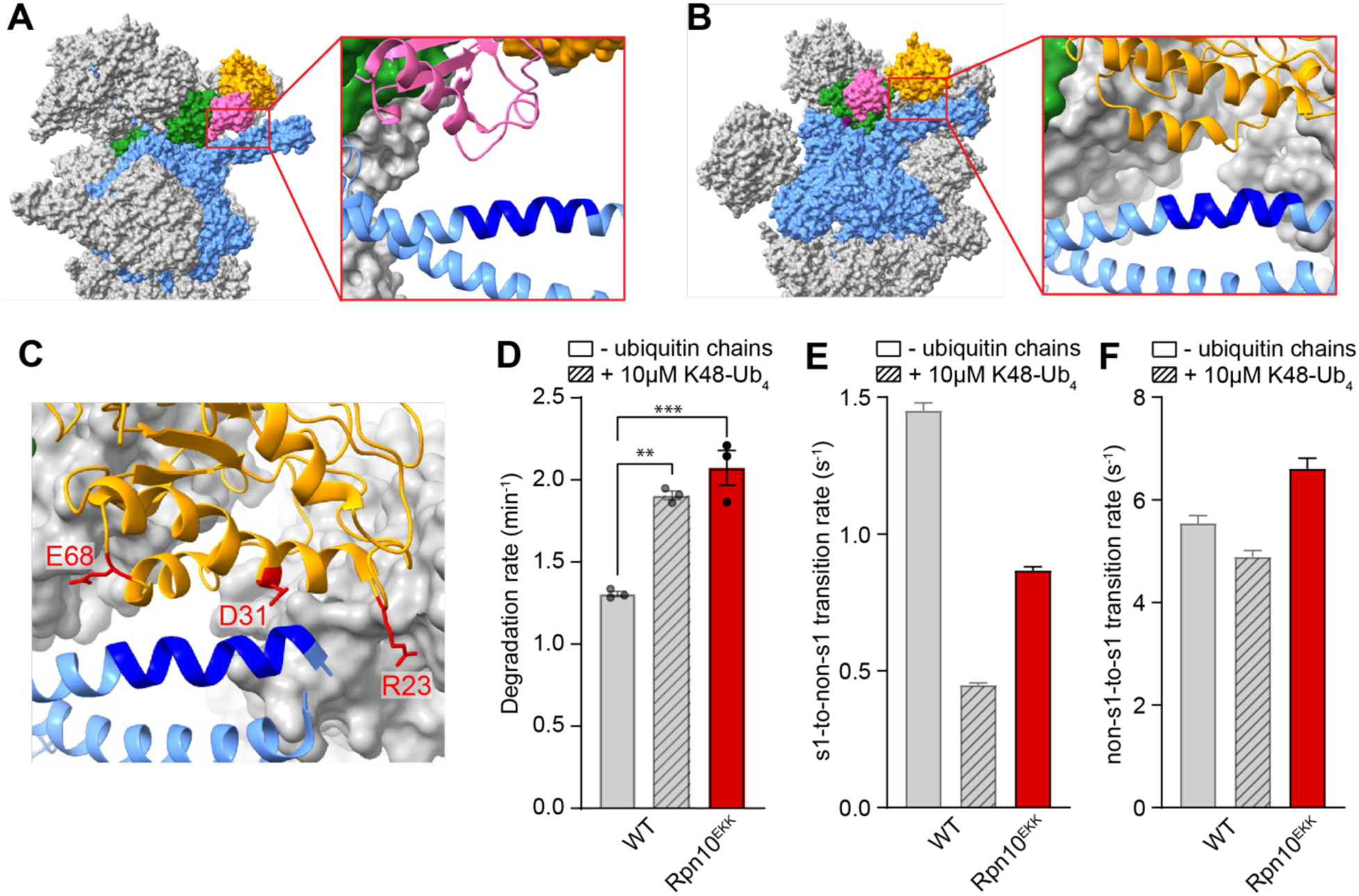
Disrupting interactions between Rpt5 and Rpn10 stabilizes the s1 state. A-B) Atomic models based on the cryo-EM structures of the engagement-competent, s1-like E_A2_ state (A, PDB ID: 6MSB) and the processing-competent, non-s1-like E_C1_ state (B, PDB ID: 6MSG) of the human 26S proteasome, with the ATPase ring shown in blue, Rpn11 in green, Rpn10’s von Willebrand factor type A (VWA) domain in orange, and ubiquitin in pink. Insets depict the details of Rpn11-bound ubiquitin contacting the same region between K51 and H64 (dark blue) of Rpt5’s coiled coil (A) as Rpn10’s VWA domain in the non-s1 state (B), leading to a competition where ubiquitin stabilizes the s1 relative to non-s1 states. C) Atomic model based on the cryo-EM structure of the *S. cerevisiae* 26S proteasome in the non-s1 state (s2 state, PDB ID: 4CR3), with Rpn10’s VWA domain contacting Rpt5’s coiled coil in a homologous region (R42-H53 shown in dark blue) as in the human proteasome (B). The critical residues R23, D31, and E68 of Rpn10 are shown in stick representation and colored red. D) Effect of the R23E, D31K, E68K triple mutations in Rpn10’s VWA domain (Rpn10^EKK^) on SspB-mediated substrate degradation. Shown are the averages of three technical replicates with error bars indicating the standard errors of the mean. Statistical significance was calculated using an ordinary one-way ANOVA test. ** p=0.0012, *** p=0.0003 E) Effect of triple mutations in Rpn10’s VWA domain (Rpn10^EKK^) on the s1-to-non-s1 transition rates as derived from the single-molecule FRET-based conformational dynamics assay. Shown are the transition rates calculated by fitting the s1-state dwell time distribution observed for at least 200 FRET-efficiency traces from two technical replicates of the proteasome conformational dynamics assay, with error bars representing the standard errors of the fit. F) Effect of triple mutations in Rpn10’s VWA domain (Rpn10^EKK^) on the non-s1-to-s1 transition rates. Shown are the transition rates calculated by fitting the s1-state dwell time distribution observed for at least 200 FRET-efficiency traces from two technical replicates of the proteasome conformational dynamics assay, with error bars representing the standard errors of the fit.

### Ubiquitin binding to Rpn11 promotes substrate engagement by stabilizing the engagement-competent state

The cryo-EM density of ubiquitin bound between Rpn11 and the Rpt5 coiled coil in the s1 state was of low resolution and highly variable among different structures ^43,44^. We were therefore unable to place specific mutations in ubiquitin that would reduce its interactions with Rpt5’s coiled-coil and confirm our hypothesis about a ubiquitin-mediated stabilization of the proteasome’s engagement-competent state. Instead, we sought to engineer a way for recruiting ubiquitin chains to bind between Rpn11 and Rpt5’s coiled-coil independent of the Rpn1, Rpn10 and Rpn13 ubiquitin receptors. We hypothesized that Rpn11 may act as the proteasome’s allosteric sensor and therefore introduced the A89F mutation, which increases Rpn11’s ubiquitin affinity by ∼ 3-fold ^45^ and turns it into a low-affinity receptor itself (Fig. 3A). Receptor-deficient proteasomes carrying this Rpn11 A89F mutation (ARR-Pru-ΔUIM-A89F) responded to the presence of K48-Ub_4_ chains with a 1.2-fold increase in SspB-delivered substrate degradation, from 1.25 min^−1^ to 1.51 min^−1^ (Fig. 3B). Consistent with our model of Rpn11 functioning as the sensor to modulate the proteasome conformational transitions, we observed in our single molecule conformational dynamics assay that ARR-Pru-ΔUIM-A89F proteasomes responded to the presence of ubiquitin chains with an attenuation in the rate of s1 to non-s1 transitions, to a similar extent as wild-type proteasomes (Fig. 3C-D).

**Fig. 3:**
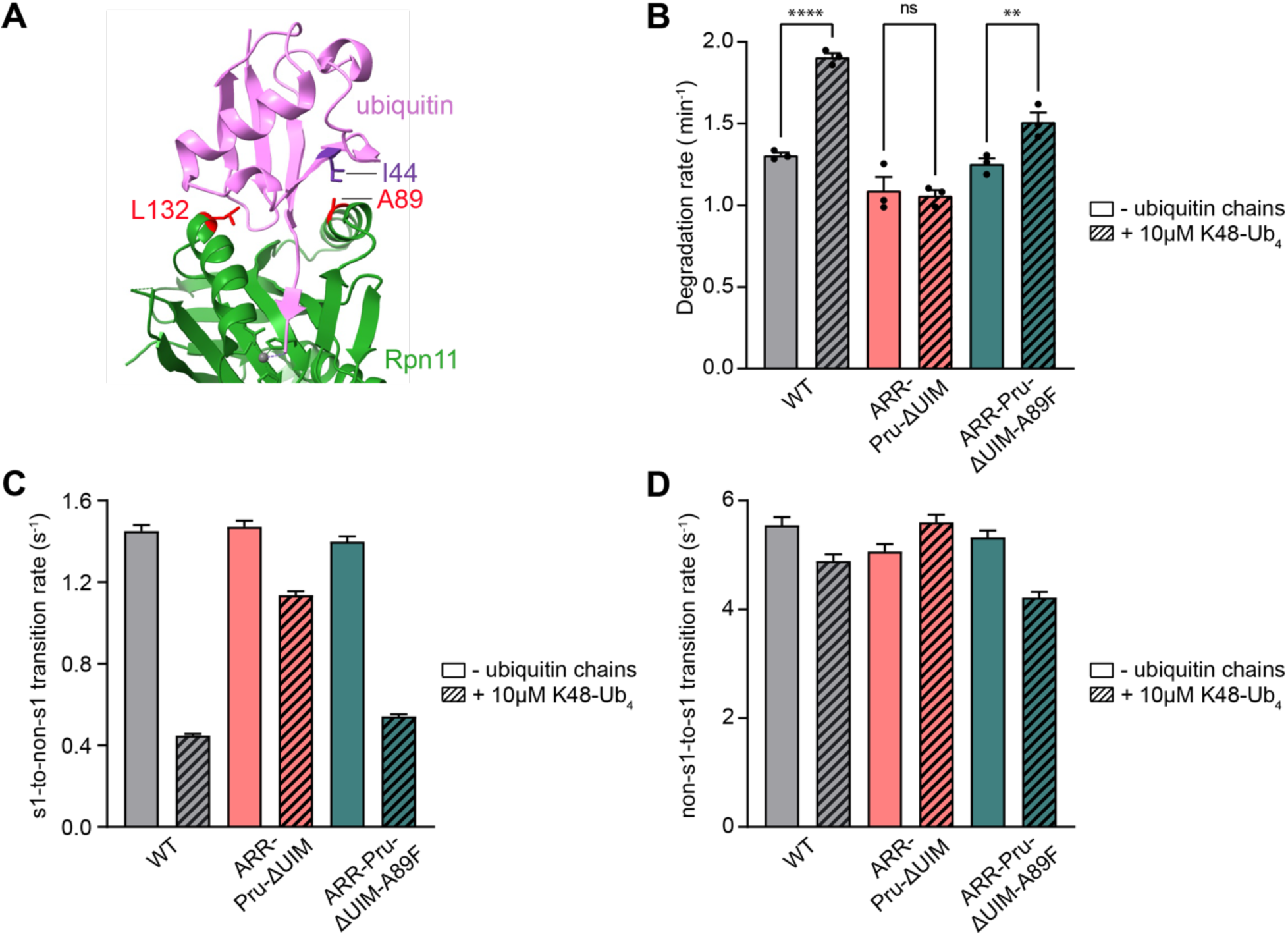
Enhancing Rpn11’s affinity for ubiquitin allows ubiquitin-mediated allosteric regulation of the proteasome in the absence of functional ubiquitin receptors. A) Interface between ubiquitin (pink) and Rpn11 (green) as seen in the crystal structure of the isolated, ubiquitin-bound Rpn11/Rpn8 heterodimer (PDB ID: 5U4P), with the critical residues A89 and L132 of Rpn11 and I44 of ubiquitin shown in stick representation. B) Effect of the Rpn11 A89F mutation on SspB-delivered substrate degradation by the triple-receptor-deficient proteasomes in the absence or presence of K48-Ub_4_ ubiquitin chains. Shown are the averages for bulk-degradation rates from three technical replicates, with error bars representing the standard error of mean. Statistical significance was calculated using an ordinary one-way ANOVA test. ns non-significant with p=0.9519, ** p=0.0071, **** p<0.0001. C-D) Effect of the Rpn11 A89F mutation on s1-to-non-s1 (C) and non-s1-to-s1 (D) transition rates of the triple-receptor-deficient proteasomes in the absence or presence of K48-Ub_4_ ubiquitin chains. Shown are the transition rates calculated by fitting the s1 and non-s1 state dwell-time distributions observed for at least 200 proteasome conformational dynamics FRET traces from two technical replicates, with error bars representing the standard errors of the fit.

We showed previously that the ubiquitin-mediated attenuation of conformational transitions and the stabilization of the engagement-competent state facilitate substrate insertion into the central channel, engagement by the ATPase motor, and consequently degradation ^36^. We therefore used our single-molecule FRET-based substrate processing assay (Fig. 4A) to confirm that ubiquitin binding to Rpn11 is responsible for the stimulatory effects of ubiquitin chains on the initial steps of substrate turnover. In this processing assay, our titin^V15P^ model substrate was labeled with the donor fluorophore LD555 on a cysteine residue within the C-terminal flexible tail, and the proteasome was labeled with the acceptor fluorophore LD655 at position 191 of the Rpt1 ATPase subunit. With this donor-acceptor pair, a successful substrate-processing event leads to a characteristic FRET-efficiency trace that starts with an intermediate value upon substrate binding to the proteasome, followed by an increase during tail insertion and engagement by the ATPase motor, a short dwell in a high-FRET state, and a subsequent gradual decay to background levels as the substrate is translocated through the channel and into the core peptidase ^36^ (Fig. 4A). In contrast, unsuccessful attempts of substrate engagement show only a short dwell in an intermediate FRET state before the substrate dissociates from the proteasome. Due to its difficult engagement and consequently too few successful degradation events, the GS substrate is not amenable for these single-molecule measurements, and we therefore used our titin^V15P^ model substrate with cyclin-B derived tail. First, we compared the effect of unanchored ubiquitin chains on the substrate capture success rate, i.e. the ratio of successful engagement events per total number of substrate encounters, for different proteasome variants. Consistent with our previous findings, the presence of ubiquitin chains doubled the capture success rate of the wild type proteasome (from 3.7 ± 0.1% to 7.8 ± 0.9 %), whereas ubiquitin did not affect proteasomes with mutated ubiquitin receptors (Fig. 4B). These data are thus also in agreement with our results for bulk substrate degradation and the proteasomal conformational switching, where the effects of ubiquitin chains depended on intact ubiquitin receptors. We therefore tested next whether enhancing Rpn11’s affinity for ubiquitin could bypass the requirement for ubiquitin receptors and stimulate the substrate-capture success of ARR-Pru-ΔUIM proteasomes. Indeed, introducing the Rpn11 A89F mutation in ARR-Pru-ΔUIM-A89F proteasome was sufficient to restore the stimulatory effect of ubiquitin chains on the substrate-capture success rate (Fig. 4B).

**Fig. 4:**
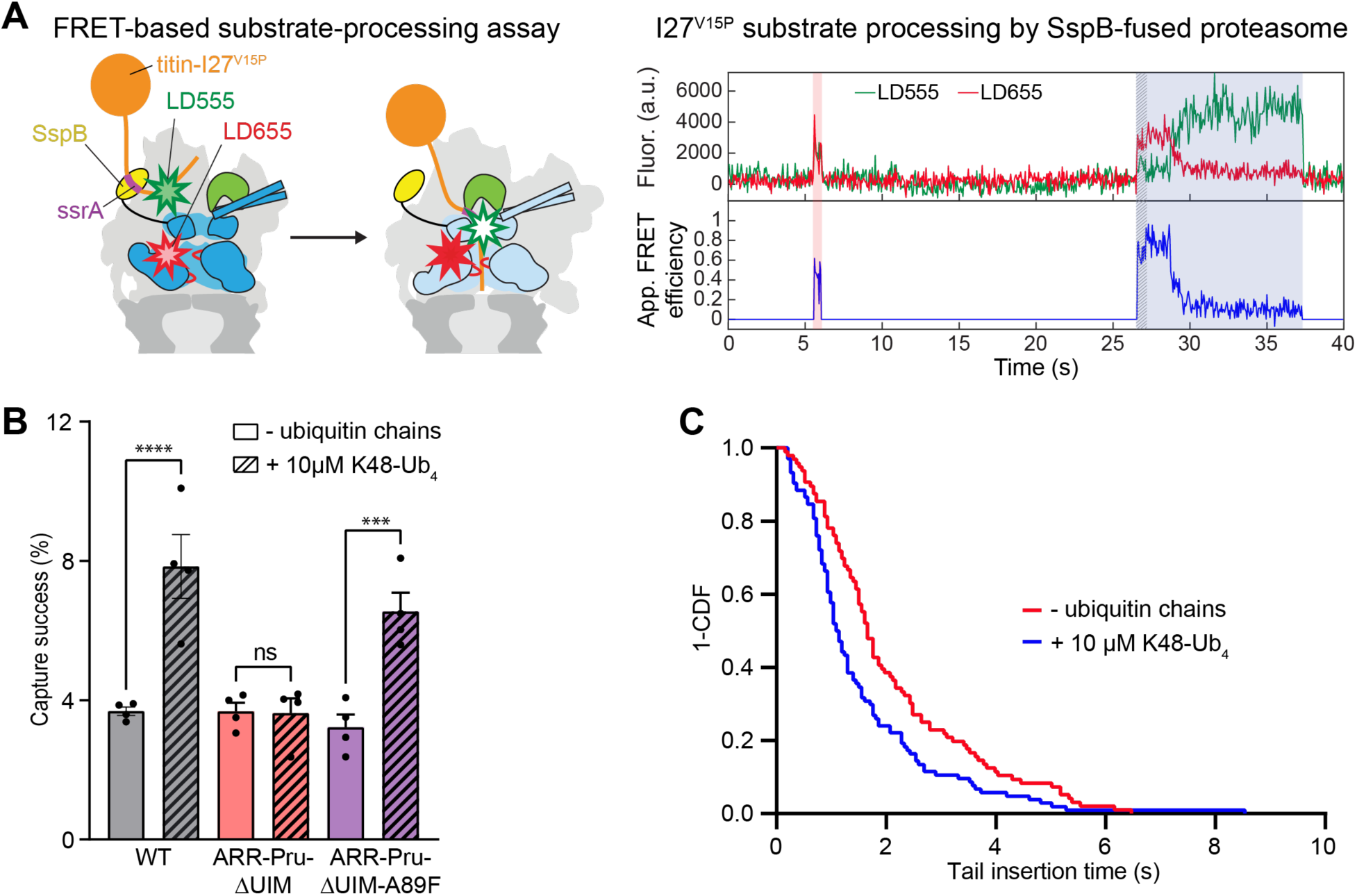
Ubiquitin binding to Rpn11 promotes substrate engagement and increases the proteasome’s substrate-capture success. A) Left: Schematic of the constructs used in the single-molecule FRET-based substrate processing assay. The ATPase motor is shown in dark blue for the engagement-competent s1 state and light blue for the substrate-engaged, processing-competent non-s1 states, Rpn11 is shown in green, the SspB fusion for substrate delivery in yellow, the titin I27^V15P^ substrate in orange, and the ssrA sequence in the substrate’s flexible initiation region for SspB binding is shown in purple. A LD555 donor dye (green star) is attached to the substrate’s initiation region and a LD655 acceptor dye (red star) is attached to the Rpt1 ATPase subunit near the central channel of the motor. Right: Example traces for the acceptor and donor fluorescence (top) and the calculated FRET efficiency (bottom) during substrate processing by an immobilized proteasome. The traces show an unsuccessful substrate-capture attempt with intermediate FRET efficiency (pink shading) and a successful capture leading to complete substrate processing (blue shading). The initial tail insertion phase is highlighted by a hatching pattern at the beginning of the substrate-processing event. B) Capture success rates for SspB-delivered substrates as determined from single-molecule processing traces for wild-type, triple-receptor-deficient (ARR-Pru-ΔUIM), and triple-receptor-deficient Rpn11^A89F^-mutant (ARR-Pru-ΔUIM-A89F) proteasomes in the absence and presence of K48-Ub_4_ ubiquitin chains. Shown are the averages from 4 technical replicates, with the error bars indicating the standard error of mean. Statistical significance was calculated using an ordinary one-way ANOVA test. ns non-significant with p=0.9998, *** p=0.0002, **** p<0.0001. D) Survival (1-CDF) plot for the tail-insertion times of SspB-delivered substrate and triple-receptor-deficient, Rp11^A89F^-mutant proteasomes (ARR-Pru-ΔUIM-A89F) in the absence (red, N = 96 events) or presence (blue, N = 104 events) of K48-Ub_4_ ubiquitin chains. Comparing the survival plot using the Gehan-Breslow-Wilcoxon test gives a p-value of 0.0005.

By analyzing the time required for the increase from intermediate to high-FRET efficiency in our substrate-processing assay, we determined the kinetics for substrate-tail insertion and engagement (τ_ins_, Fig. 4A). Unanchored K48-Ub_4_ chains decreased the average tail-insertion time constant for the wildtype proteasome from 1.64 ± 0.10 s to 1.39 ± 0.10 s (Supplementary Fig. 5A, 6A, B, Supplementary Table 2). As for the other allosteric effects described above, this ubiquitin-mediated acceleration of tail insertion depended on functional ubiquitin receptors, as no statistically significant acceleration was observed for ARR-Pru-ΔUIM proteasomes (Supplementary Fig. 5, 6 B-D, supplementary Table 2), but the effects were restored upon introducing the Rpn11 A89F mutation in ARR-Pru-ΔUIM-A89F proteasomes (Fig. 4C, Supplementary Fig. 6E,F, Supplementary Table 2). Our findings that the Rpn11 A89F mutation can compensate for the lack of ubiquitin receptors in mediating the allosteric effects of ubiquitin chains on the proteasome conformational switching, substrate-tail insertion, and capture success indicates that Rpn11 is indeed the proteasome’s main allosteric sensor for ubiquitin.

To test this further, we returned to the more sensitive, slowly engaging GS substrate and modified its flexible initiation region with just a mono-ubiquitin (supplementary Fig. 7A), which is unable to simultaneously contact a ubiquitin receptor and Rpn11, but should be sufficient to allosterically trigger Rpn11 in our SspB-mediated substrate delivery system. Indeed, the degradation of mono-ubiquitinated GS substrate by the SspB-fused ARR-Pru-ΔUIM proteasome was two-fold faster than the degradation of the non-ubiquitinated GS substrate (0.38 ± 0.02 min^−1^ versus 0.21 ± 0.02 min^−1^; Fig. 5A, supplementary Fig. 7B). For additional validation we characterized two Rpn11 mutations that increase the affinity for ubiquitin, A89F and A89I, and one mutation designed to attenuate ubiquitin binding, L132R (Fig. 3A). For the isolated Rpn11 outside the proteasome context, the A89F and A89I mutations stimulated the deubiquitinase activity 1.3-fold and 10-fold, respectively, while the L132R mutation eliminated deubiquitination (Fig. 5B, supplementary Fig. 7C). In the context of SspB-mediated delivery of a mono-ubiquitinated ssrA-tagged substrate, we did not expect a considerable effect for the affinity-increasing A89F and A89I mutations, as the SspB-ssrA interaction leads to an artificially high local ubiquitin concentration near Rpn11. Consistently, SspB-fused ARR-Pru-ΔUIM proteasomes with Rpn11^A89F^ or Rpn11^A89I^ variants showed wild-type-like degradation kinetics, with degradation rates of 0.42 ± 0.02 min^−1^ and 0.38 ± 0.01 min^−1^, respectively, for the mono-ubiquitinated GS substrate, and degradation rates of 0.21 ± 0.005 min^−1^ and 0.22 ± 0.05 min^−1^, respectively, for the non-ubiquitinated substrate (Fig. 5C, supplementary Fig. 7D,E). In contrast, SspB-fused ARR-Pru-ΔUIM proteasomes containing the deleterious Rpn11^L132R^ mutation showed no ubiquitin-mediated stimulation of GS substrate degradation, with degradation rates of 0.25 ± 0.02 min^−1^ and 0.23 ± 0.02 min^−1^ for the mono-ubiquitinated and non-ubiquitinated substrates, respectively (Fig. 5C, supplementary Fig. 7F). Even at local high ubiquitin concentrations, the L132R mutation thus prevents a functional Rpn11 binding.

**Figure 5:**
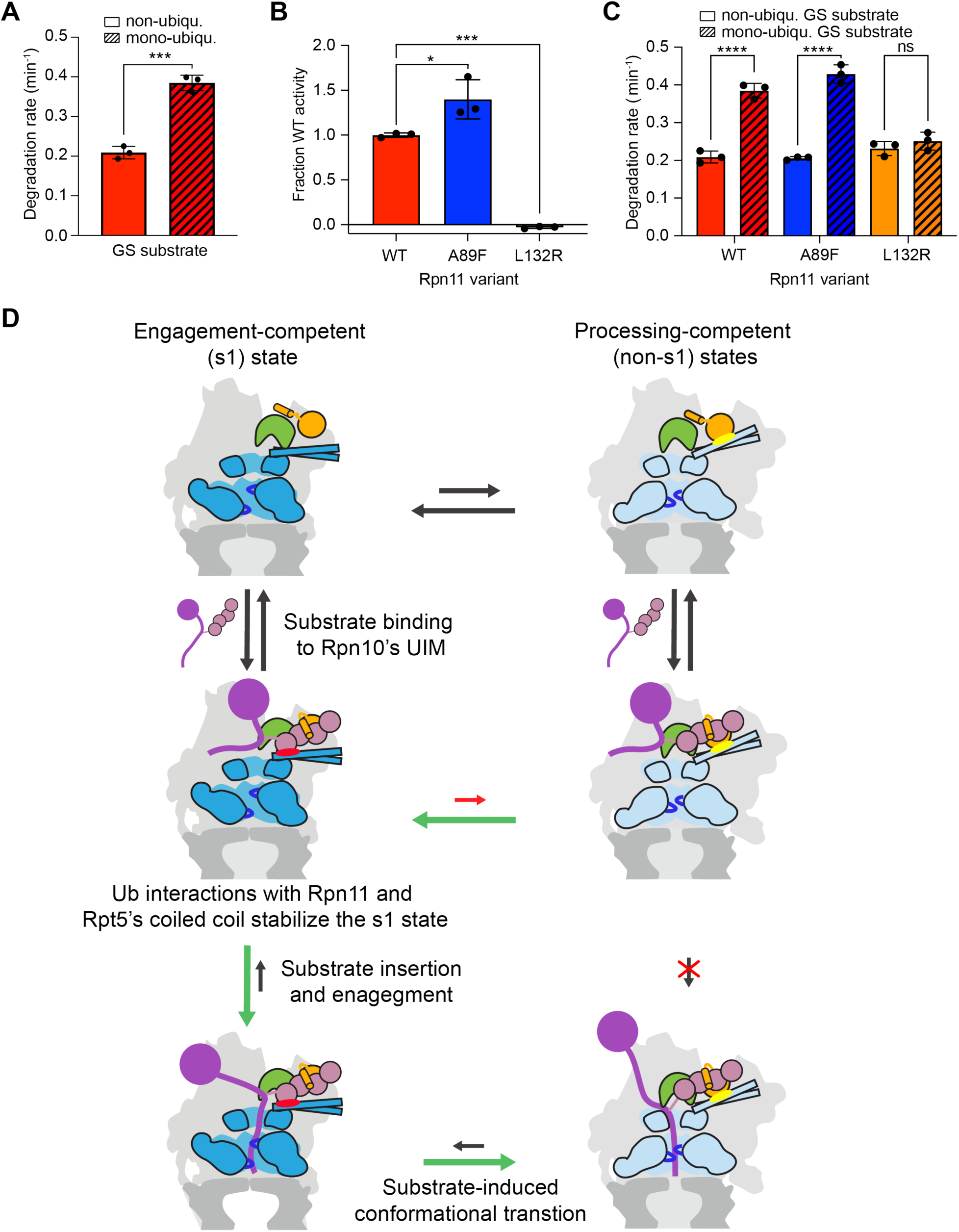
Rpn11 functions as the allosteric ubiquitin sensor. A) Rates for the SspB-mediated degradation of the slowly engaging GS substrate in its unmodified or mono-ubiquitinated form by triple-receptor-deficient 26S proteasome. Shown are the averages of three technical replicates with error bars indicating the standard deviation. Statistical significance was calculated using a standard unpaired t-test *** p = 0.0003. B) Deubiquitinase activity of the isolated Rpn11/Rpn8 heterodimer containing the Rpn11 A89F or L132R mutation at the ubiquitin-binding interface. Shown are the averages of three technical replicates with error with error bars representing standard deviation. Statistical significance was calculated using an ordinary one-way ANOVA test: * p = 0.0151, *** p = 0.001. C) Rates for the SspB-mediated degradation of the slowly engaging GS substrate in its unmodified or mono-ubiquitinated form by triple-receptor-deficient 26S proteasome with wild-type Rpn11, Rpn11 A89F, or Rpn11 L132R. Shown are the averages of three technical replicates with error bars indicating the standard deviation. Statistical significance was calculated using an ordinary one-way ANOVA test: **** p<0.0001, ns not significant with p=0.8005. D) Model for the allosteric regulation of proteasomal substrate engagement through early ubiquitin binding to Rpn11. The substrate-free proteasome spontaneously switches between the engagement-competent s1 state (left) and the processing-competent non-s1 states (right). Binding of a ubiquitinated substrate (purple with pink ubiquitin chain) to the Rpn10 receptor (orange) allows ubiquitin binding to Rpn11 (green) and a simultaneous contact with Rpt5’s coiled coil (red ellipse), which stabilizes the s1 state by competing with the non-s1-state specific interaction between Rpn10’s VWA domain and Rpt5’s coiled coil (yellow ellipse). This shifts the conformational equilibrium towards the s1 state with an accessible central channel and thus facilitates the insertion of a substrate’s flexible initiation region into the ATPase motor. Upon successful substrate engagement, the proteasome switches to non-s1 processing-competent states for processive substrate translocation, unfolding, and co-translocational deubiquitination.

Combined, these data indicate that Rpn11 acts as an allosteric sensor whose interaction with ubiquitin, prior to the co-translocational deubiquitination, attenuates the conformational switching of the proteasome and thus facilitates the engagement of ubiquitin-attached polypeptide substrates by the ATPase motor.

## Discussion

Our biochemical and single-molecule studies elucidate the mechanisms by which ubiquitin chains allosterically regulate the proteasome conformational switching and the engagement of a substrate’s flexible initiation region for degradation. Intriguingly, the allosteric sensor for this regulation is the main proteasomal deubiquitinase Rpn11, whose initial binding of ubiquitin allows this ubiquitin moiety to form a stabilizing interaction with the Rpt5 coiled coil and increase the lifetime of the proteasome’s engagement-competent s1 state for substrate insertion into the central channel. We propose a model (Fig. 5D) where an ubiquitinated substrate is recruited to the proteasome through interactions with Rpn10’s UIM, which allows ubiquitin to immediately bind the nearby Rpn11 in a pre-cleavage state, simultaneously contact the Rpt5 coiled coil, and thereby prevent the spontaneous conformational transition of the proteasome to the processing-competent non-s1 states with an occluded channel entrance. We previously found that Rpn11’s Insert-1 region forms an inhibitory loop over the catalytic groove and that the transition of this loop to an active-state hairpin represents the rate-limiting step for ubiquitin cleavage ^24^. Mechanical pulling on an engaged ubiquitinated substrate by the ATPase motor accelerates the loop-to-hairpin transition in the proteasome-incorporated Rpn11 and allows co-translocational deubiquitination with a time constant of ∼ 1 s ^36^, whereas in the absence of substrate translocation, for instance with an engagement-incompetent substrate, deubiquitination occurs much more slowly with a time constant of ∼ 45 s ^12^. The inactive Insert-1 loop conformation is thus long-lived enough to prevent any premature deubiquitination prior to substrate engagement, while the Rpn11-bound ubiquitin acts as an allosteric regulator and facilitates substrate insertion into the ATPase motor. Upon successful motor engagement of the substrate’s flexible initiation region, the proteasome switches to the processing-competent non-s1 states ^12,36^ and mechanical pulling on the substrate accelerates the loop-to-hairpin transition of Rpn11’s Insert-1 region for rapid co-translocational deubiquitination ^24^. That ubiquitin binds to Rpn11 for allosteric regulation right after substrate landing on the proteasome and prior to substrate insertion into the ATPase motor is also supported by our previous findings for the Rpn11 G77P mutation, which favors the active state hairpin conformation of Rpn11, leads to faster and thus premature deubiquitination, and causes substrate release from the proteasome ^24^.

We observed that extending the lifetime of the proteasome’s engagement-competent s1 state through ubiquitin binding to Rpn11 has a moderate effect on the degradation of our model substrate, which contains an ideal, 35-residue C-terminal initiation region and an optimally placed ubiquitin chain. However, most cellular substrates are likely more challenging to grab by the proteasome, for instance due to non-ideal ubiquitin-chain placement and length, or a slowly engageable, short, or slippery initiation region. The degradation of those substrates is expected to significantly benefit from an ubiquitin- and Rpn11-medited s1-lifetime extension, which facilitates substrate insertion, thereby increases the capture success, and accelerates overall degradation velocity, similar to our GS substrate that showed ∼ 4-fold faster turnover in the presence of ubiquitin ^36^. Our findings also help to advance our understanding of how the geometry of substrate delivery, the position of ubiquitin, or the type of an ubiquitin modification, i.e. the length, linkage, and branching of a chain, could affect degradation. There have been conflicting reports about the dependence of different substrates on proteasomal ubiquitin receptors, in particular Rpn10’s UIM ^23,38,46^, and our model may explain the mechanistic basis for these discrepancies. Depending on the length and sequence of a substrate’s initiation region as well as the position and/or length of a ubiquitin chain, the Rpn11/ubiquitin-mediated attenuation of the proteasome conformational switching could become critical for successful substrate engagement, and substrate delivery through Rpn10’s UIM could therefore be more favorable compared to the Rpn1 or Rpn13 receptors. This also means that the efficiency of PROteolysis TArgeting Chimeras (PROTACs) can potentially be optimized by finetuning the ubiquitination position and delivery geometry of a targeted neo-substrate. Future work will be required to elucidate the complex relationship between the substrate-attached ubiquitin code and proteasomal ubiquitin receptors in regulating protein turnover, as well as the enigmatic roles of various regulatory proteins such as ubiquitin ligases and shuttle receptors that transiently associate with the proteasome. Several E3 ubiquitin ligases, including Ube3A in higher eukaryotes ^47^, Ubr1 and Ufd4 in budding yeast ^48^, and Upl3 in plants ^49^, have been proposed to extend ubiquitin modifications on proteasome-targeted substrates. One attractive model is that these ligase activities adjust substrate ubiquitination patterns or delivery geometries to facilitate the ubiquitin-mediated promotion of substrate engagement. In yeast, the shuttle factors Rad23 and Dsk2 have been proposed to serve as alternative ubiquitin receptors for the delivery of some proteasomal substrates ^50^. Interestingly, Rad23 and Dsk2 preferentially bind to Rpn1 and Rpn13, respectively ^51^ and it is possible that they assist ubiquitin chains and Rpn11 in sterically attenuating the proteasome conformational dynamics, which will be exciting areas to investigate in the future.

In summary, our study reveals the mechanism by which the Rpn11 deubiquitinase fulfills a second function, as an allosteric ubiquitin sensor in promoting substrate engagement by the proteasomal ATPase motor prior to deubiquitination, and thus leads to a significant advantage for the turnover of ubiquitinated versus non-ubiquitinated proteins in a crowded cellular environment. Our model presents intriguing new ideas for investigating the role of different ubiquitin receptors, shuttle factors, and transiently-associated ubiquitin ligases in regulating proteasomal substrate degradation.

## Acknowledgements

We thank the members of the Martin lab for helpful discussions.

## Author Contributions

Conceptualization, Z.M.H. and A.M.; Methodology, Z.M.H. and A.M.; Investigation, Z.M.H. and K.C.D.; Writing – Original Draft, Z.M.H. and A.M.; Writing – Review & Editing, Z.M.H., K.C.D., and A.M.; Funding Acquisition, A.M.; Supervision, A.M.

## Supplementary Figures and Tables

**Supplementary Fig. 1:**
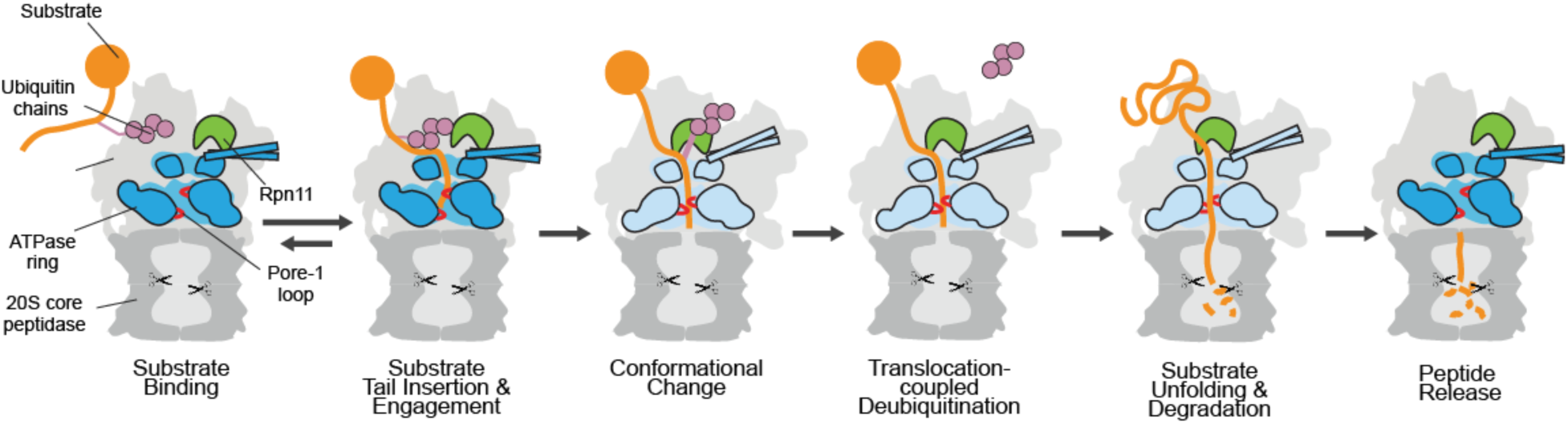
Current model for individual steps of substrate degradation by the 26S proteasome. The cutaway view of the 26S proteasome shows the 20S core particle in dark grey, the lid and non-ATPase subunits of the base subcomplex in light grey, the ATPase hexamer in blue, translocating pore-1 loops in red, the Rpn11 deubiquitinase in green, the substrate in orange, and the substrate-attached poly-ubiquitin chain in purple. After a substrate is recruited through ubiquitin binding to a proteasomal receptor, the substrate’s flexible initiation region diffuses into the central channel of the ATPase ring. Upon successful substrate engagement with the pore-1 loops, the base ATPase transitions from the engagement-competent s1 state (dark blue) to processing-competent non-s1 states (light blue) for substrate unfolding and processive translocation into the core particle for proteolytic cleavage. Ubiquitin chains are removed from the substrate by Rpn11 in a co-translocational manner. After complete threading of the substrate polypeptide, the base ATPase switches back to the engagement-competent s1 state.

**Supplementary Fig. 2:**
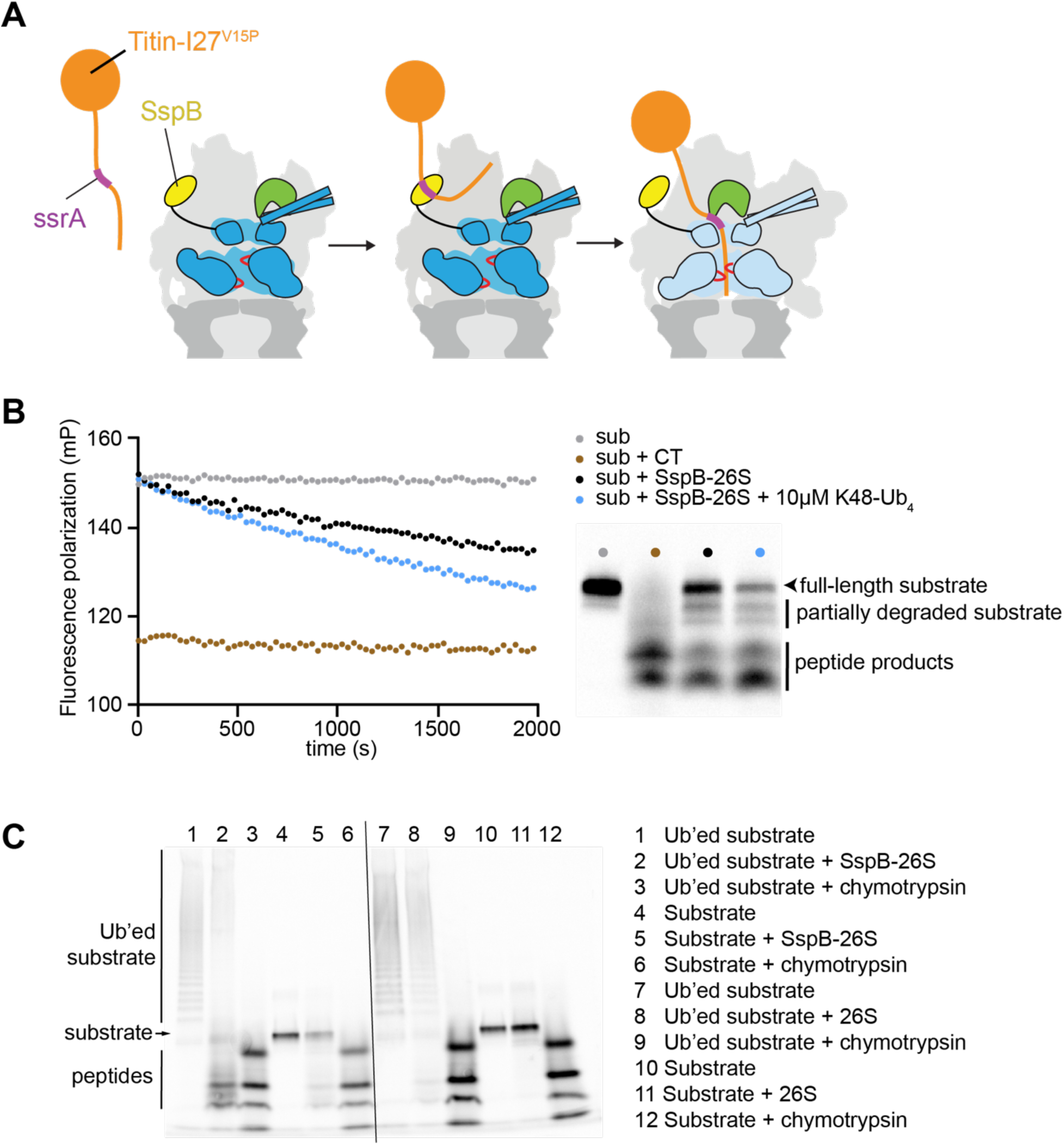
Ubiquitin-independent SspB-mediated substrate degradation by the 26S proteasome. A) Schematic for the degradation of a titin^V15P^ model substrate (orange) with the ssrA degron sequence (purple) in the C-terminal unstructured initiation region. The engineered proteasome variant contains the ssrA-binding SspB adaptor (yellow) from *E. coli* fused to the N-terminus of the Rpt2 ATPase subunit of the base subcomplex (blue). B) Left: Example traces for the degradation of the ssrA-tagged titin^V15P^ model substrate by SspB-fused 26S proteasome in the absence or presence of unanchored K48-linked tetra-ubiquitin chains (K48-Ub_4_). The substrate was N-terminally labeled with fluoresceine amidite (FAM) to monitor degradation by fluorescence polarization. Also shown is the complete cleavage by chymotrypsin as a control. Right: SDS-PAGE analysis of aliquots taken after 60 min from the polarization- monitored degradation samples. C) SDS-PAGE (4 - 20%) analysis of the endpoints for the degradation of ubiquitinated (Ub’ed) or unmodified ssrA-containing titin^V15P^ substrate by SspB-fused or wild-type 26S proteasome or chymotrypsin.

**Supplementary Fig. 3:**
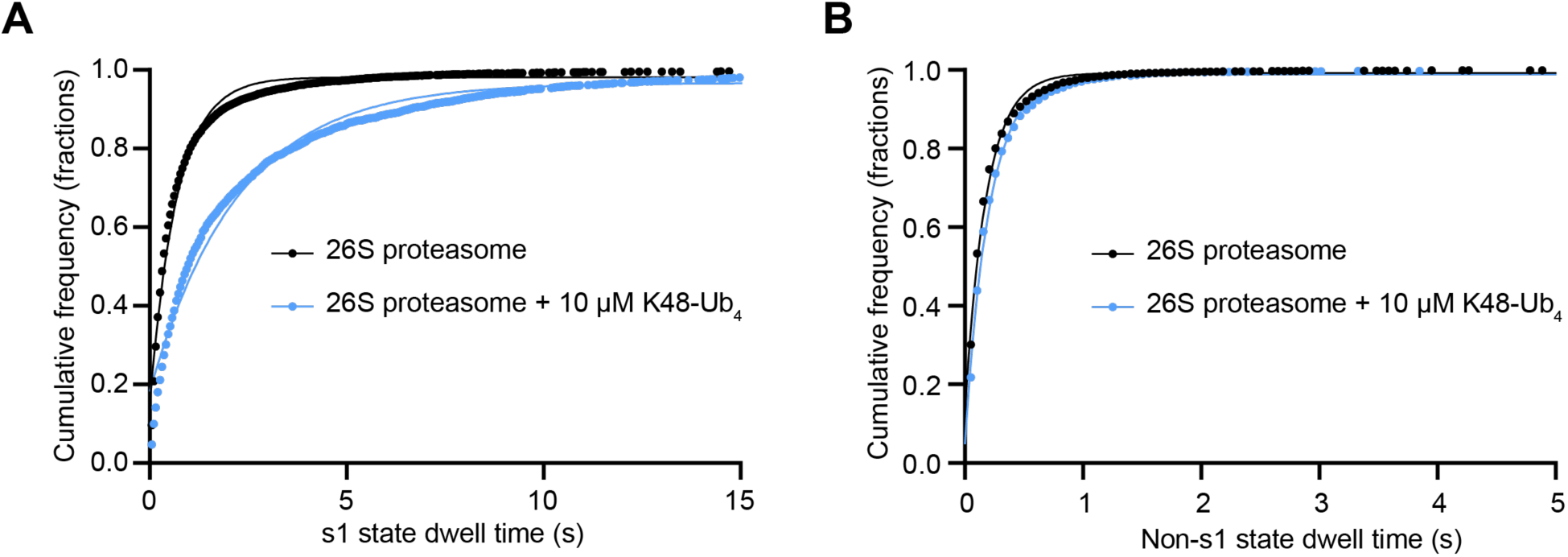
Determination of the transition rates of the proteasome conformations. Cumulative frequencies for the dwell-time distributions of the low-FRET s1 state (A) and the high-FRET non-s1 states (B) during the conformational switching of the substrate-free wild-type proteasome in the absence of K48-Ub_4_ (black, N = 227 events for s1 state, N = 73 events for non-s1 states) and in the presence of K48-Ub_4_ (blue, N = 295 events for s1 state, N = 53 events for non-s1 states), with fits to single exponentials shown as black and blue lines.

**Supplementary Fig. 4:**
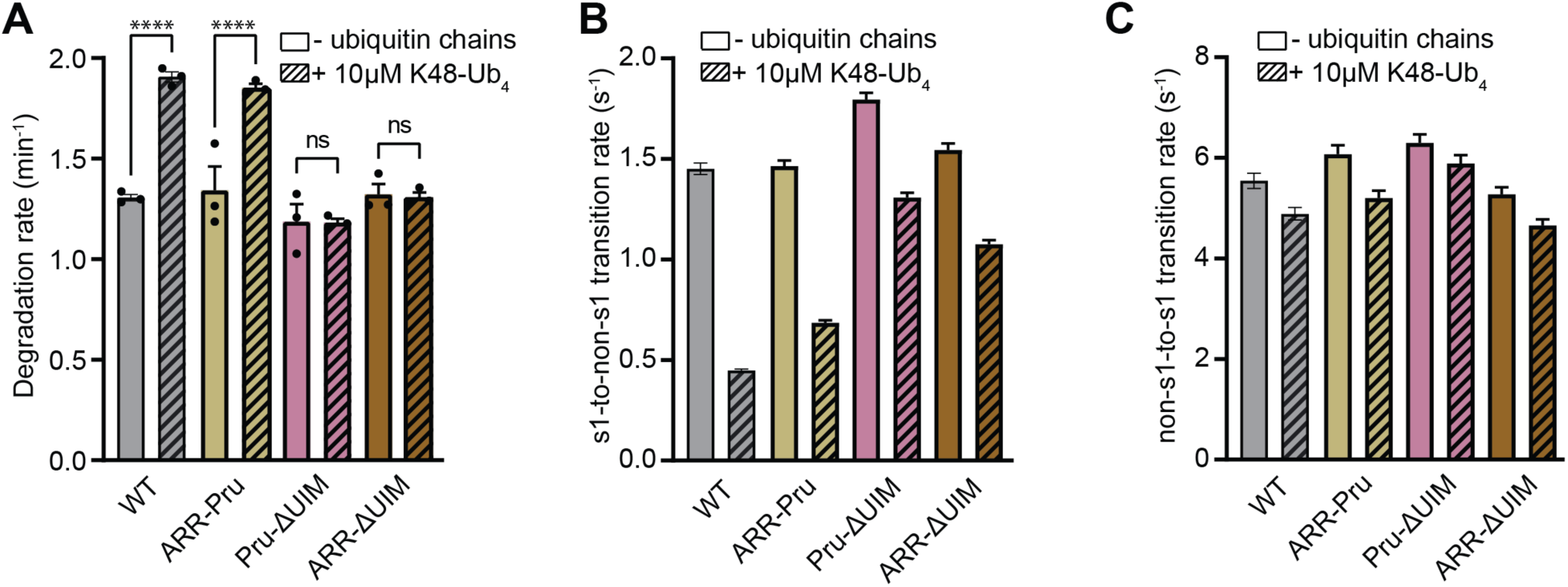
The ubiquitin interacting motif of Rpn10 mediates the allosteric effect of ubiquitin chains on the proteasomal substrate degradation and conformational dynamics. A) Unanchored K48-Ub_4_ ubiquitin-chain dependent stimulation of SspB-delivered substrates degradation by wild-type and receptor-deficient 26S proteasomes, carrying a combination of mutations in the T1 site of Rpn1 (ARR), mutations in the Pru domain of Rpn13 (Pru), or a deletion of Rpn10’s UIM (ΔUIM). Shown are the averages of three technical replicates with error bars representing the standard errors of mean. Statistical significance was calculated using an ordinary one-way ANOVA test. ns p>0.9999, **** p<0.0001. B) Effects of unanchored K48-Ub_4_ on the s1-to-non-s1-transition rates for wild-type and receptor-deficient proteasomes with a combination of ARR, Pru, or ΔUIM mutations. Shown are the transition rates calculated by fitting the s1-state dwell time distribution observed in at least 200 FRET-efficiency traces from two technical replicates of the proteasome conformational dynamics assay, with error bars representing the standard errors of the fit. C) Effects of unanchored K48-Ub_4_ on the non-s1-to-s1 transition rates for wild-type and receptor-deficient proteasomes with a combination of ARR, Pru, or ΔUIM mutations Shown are the transition rates calculated by fitting the non-s1-state dwell time distribution observed in at least 200 FRET-efficiency traces from two technical replicates of the proteasome conformational dynamics assay, with error bars representing the standard errors of the fit.

**Supplementary Fig. 5:**
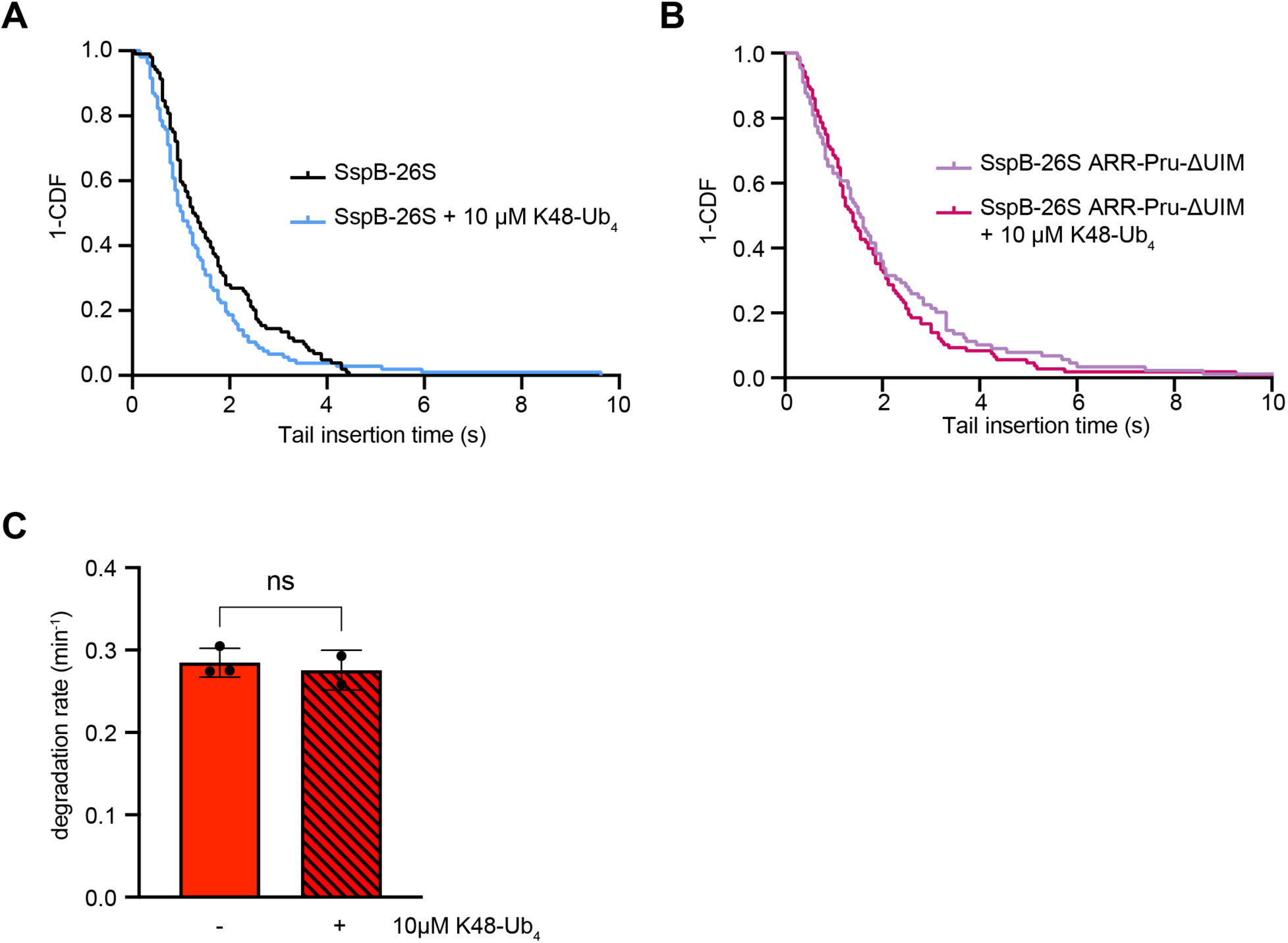
Effects of ubiquitin chains on substrate tail insertion and degradation kinetics. A,B) Survival (1-CDF) plots for the tail-insertion times of SspB-delivered titin^V15P^ substrate and wild-type (A) or triple-receptor-deficient, ARR-Pru-ΔUIM-mutant proteasome (B) in the absence (black, N = 104 events; purple, N = 89 events) or the presence (blue, N = 107 events; magenta, N = 108 events) of K48-Ub_4_ ubiquitin chains. Comparing the survival plot using the Gehan-Breslow-Wilcoxon test gives p-values of 0.0189 and 0.7842 for wild-type and triple-receptor-deficient, ARR-Pru-ΔUIM-mutant proteasome, respectively. C) Rates for the degradation of SspB-delivered GS substrate triple-receptor-deficient (ARR-Pru-ΔUIM) 26S proteasome in the absence and presence of K48-Ub_4_ ubiquitin chains. Shown are the averages from 3 technical replicates, with the error bars indicating the standard error of mean. Statistical significance was calculated using an ordinary one-way ANOVA test. ns non-significant with p = 0.6526.

**Supplementary Fig. 6:**
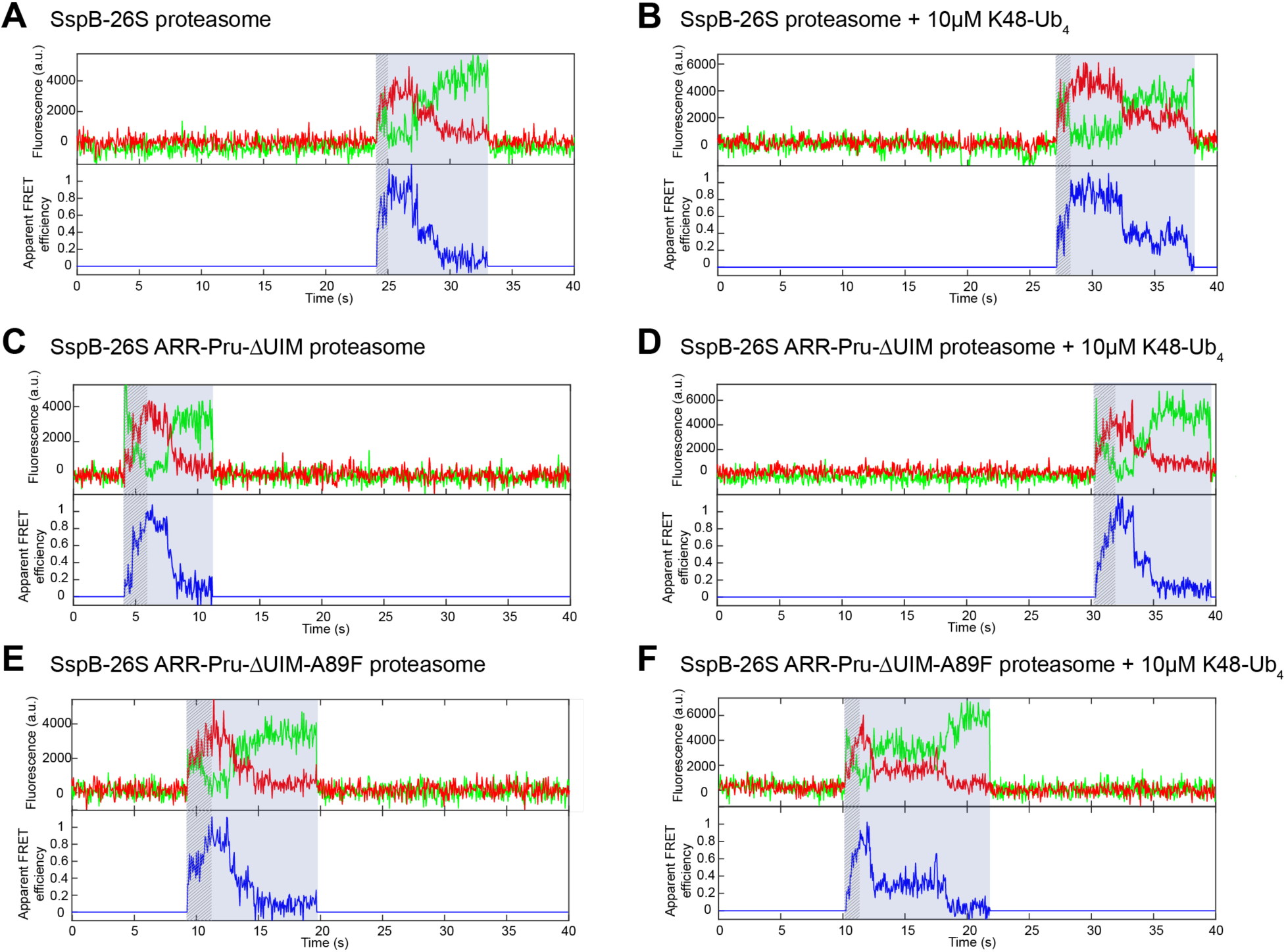
Representative traces for the single-molecule substrate-processing assay monitoring the degradation of the SspB-delivered titinV15P model substrate. Substrate degradation by SspB-fused wild-type proteasome in the absence (A) and presence (B) of unanchored tetraubiquitin chains K48-Ub_4_. Substrate degradation by SspB-fused triple-receptor-deficient (ARR-Pru-ΔUIM) proteasome in the absence (C) and presence (D) of unanchored tetraubiquitin chains K48-Ub_4_. Substrate degradation by SspB-fused triple-receptor-deficient proteasome with Rpn11 A89F mutation ((ARR-Pru-ΔUIM-A89F) in the absence (E) and presence (F) of unanchored tetraubiquitin chains K48-Ub_4_.

**Supplementary Fig. 7:**
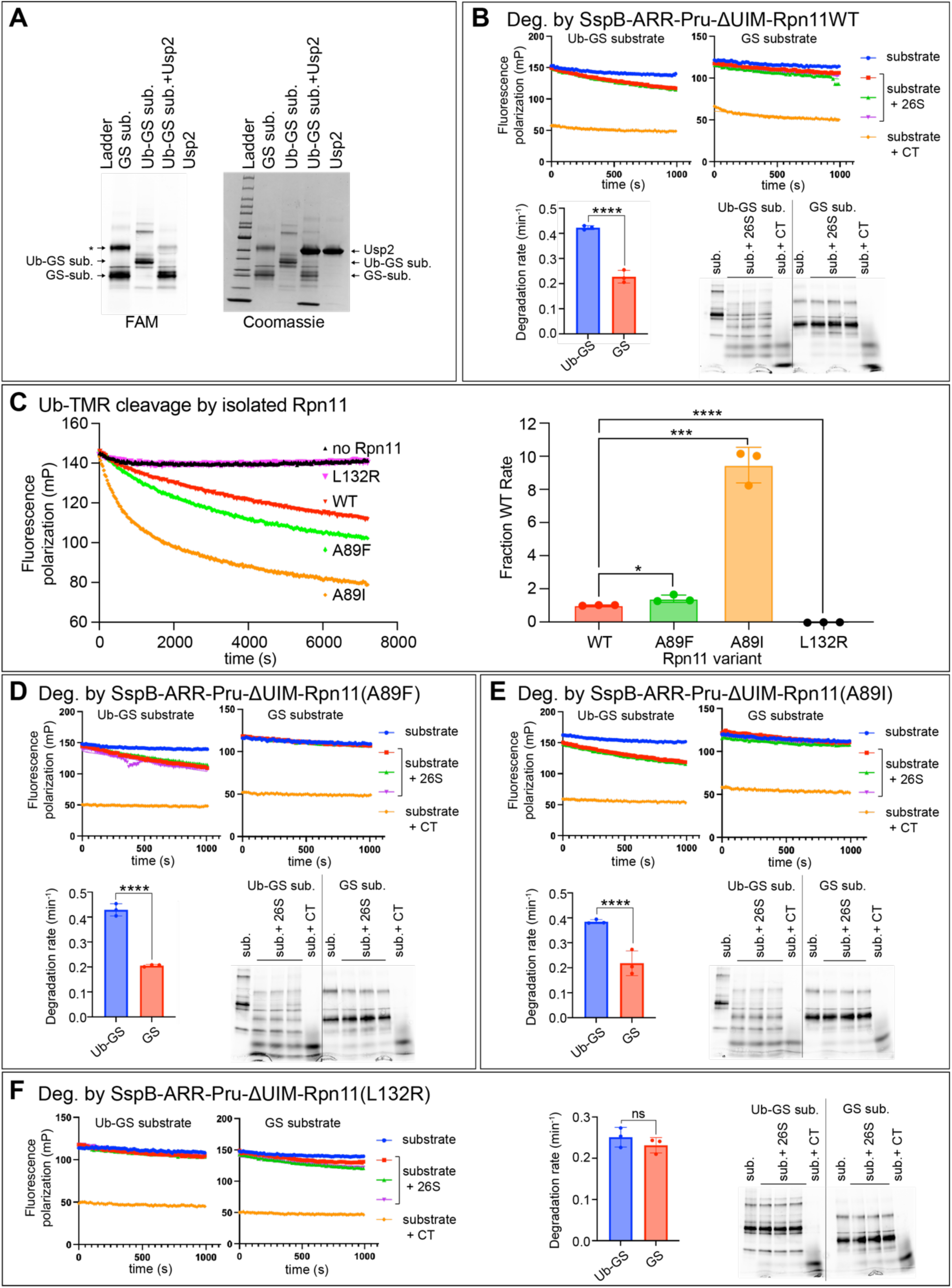
Characterization of Rpn11-mutant effects on de-ubiquitination activity and GS-substrate degradation. A) Fluorescence scan (left) and Coomassie stain of a 4-20% SDS-PAGE gel showing the fluoresceine (FAM) labeled non-ubiquitinated GS-substrate and ubiquitinated Ub-GS-substrate before and after treatment with the deubiquitinase Usp2. * marks a dimeric artifact of the GS-substrate. B) SspB/ssrA-mediated degradation of the GS- substrate and Ub-GS substrate by the ubiquitin-receptor-less, SspB-fused 26S proteasome. Top: Example traces for the fluorescence polarization changes of the Ub-GS and GS substrates alone, during multiple-turnover proteasomal degradation, or in the presence of chymotrypsin (CT). Bottom left: calculated degradation rates. Shown are the averages of three technical replicates with error bars representing the standard errors of the mean. Statistical significance was calculated using an ordinary one-way ANOVA test. **** p < 0.0001. Bottom right: Coomassie-stained SDS-PAGE (4-20%) analysis of the degradation end points. C) Cleavage of Ubiquitin-TAMRA by isolated Rpn11/Rpn8 and its variants. Left: representative fluorescence-polarization traces for the cleavage of Ub-TAMRA by wild-type Rpn11 and its mutants. Right: Calculated relative rates for the deubiquitination activities of wild-type and mutant Rpn11. Shown are the averages of three technical replicates with error bars representing the standard errors of the mean. Statistical significance was calculated using an unpaired t-test. * p = 0.0348, *** p = 0.0002, **** p <0.0001. D) SspB/ssrA-mediated degradation of the GS-substrate and Ub-GS substrate by the ubiquitin-receptor-less, SspB-fused 26S proteasome with Rpn11 A89F mutation. Top: Example traces for the fluorescence polarization changes of the Ub-GS and GS substrates alone, during multiple-turnover proteasomal degradation, or in the presence of chymotrypsin (CT). Bottom left: calculated degradation rates. Shown are the averages of three technical replicates with error bars representing the standard errors of the mean. Statistical significance was calculated using an ordinary one-way ANOVA test. **** p < 0.0001. Bottom right: Coomassie-stained SDS-PAGE (4-20%) analysis of the degradation end points. E) SspB/ssrA-mediated degradation of the GS-substrate and Ub-GS substrate by the ubiquitin-receptor-less, SspB-fused 26S proteasome with Rpn11 A89I mutation. Top: Example traces for the fluorescence polarization changes of the Ub-GS and GS substrates alone, during multiple-turnover proteasomal degradation, or in the presence of chymotrypsin (CT). Bottom left: calculated degradation rates. Shown are the averages of three technical replicates with error bars representing the standard errors of the mean. Statistical significance was calculated using an ordinary one-way ANOVA test. **** p < 0.0001. Bottom right: Coomassie-stained SDS-PAGE (4- 20%) analysis of the degradation end points. F) SspB/ssrA-mediated degradation of the GS- substrate and Ub-GS substrate by the ubiquitin-receptor-less, SspB-fused 26S proteasome with Rpn11 L132R mutation. Left: Example traces for the fluorescence polarization changes of the Ub-GS and GS substrates alone, during multiple-turnover proteasomal degradation, or in the presence of chymotrypsin (CT). Right: calculated degradation rates and Coomassie-stained SDS-PAGE (4-20%) analysis of the degradation end points. For the degradation rates, shown are the averages of three technical replicates with error bars representing the standard errors of the mean. Statistical significance was calculated using an ordinary one-way ANOVA test. ns non-significant with p = 0.8005.

**Supplementary Table 1:** Rates of proteasome conformational switching. Rates for the conformational switching from the s1 to non-s1 states (k_s1_) and from non-s1 to s1 states (k_non-s1_) were determined by Hidden-Markov-modeling of the results from the FRET-based conformational dynamic assay for wild-type proteasome and various proteasome variants in the absence and presence of unanchored K48-linked tetraubiquitin chains.

| **Proteasome variant** | **$k_{s1}$ ($s^{-1}$)** | **$k_{non-s1}$ ($s^{-1}$)** |
| --- | --- | --- |
| WT proteasome | $1.45 \pm 0.03$ | $5.54 \pm 0.15$ |
| WT proteasome + 10 μM K48-Ub<sub>4</sub> | $0.45 \pm 0.01$ | $4.89 \pm 0.13$ |
| ΔUIM proteasome | $1.64 \pm 0.03$ | $5.54 \pm 0.15$ |
| ΔUIM proteasome + 10 μM K48-Ub<sub>4</sub> | $1.29 \pm 0.03$ | $5.01 \pm 0.11$ |
| ARR proteasome | $1.28 \pm 0.03$ | $5.07 \pm 0.14$ |
| ARR proteasome + 10 μM K48-Ub<sub>4</sub> | $0.58 \pm 0.01$ | $4.16 \pm 0.10$ |
| Pru proteasome | $1.67 \pm 0.03$ | $6.44 \pm 0.17$ |
| Pru proteasome + 10 μM K48-Ub<sub>4</sub> | $0.62 \pm 0.01$ | $5.13 \pm 0.14$ |
| ARR-Pru proteasome | $1.46 \pm 0.03$ | $6.08 \pm 0.18$ |
| ARR-Pru proteasome + 10 μM K48-Ub<sub>4</sub> | $0.69 \pm 0.01$ | $5.20 \pm 0.15$ |
| Pru-ΔUIM proteasome | $1.80 \pm 0.03$ | $6.30 \pm 0.16$ |
| Pru-ΔUIM proteasome + 10 μM K48-Ub<sub>4</sub> | $1.31 \pm 0.02$ | $5.89 \pm 0.17$ |
| ARR-ΔUIM proteasome | $1.54 \pm 0.03$ | $5.27 \pm 0.14$ |
| ARR-ΔUIM proteasome + 10 μM K48-Ub<sub>4</sub> | $1.08 \pm 0.02$ | $4.65 \pm 0.12$ |
| ARR-Pru-ΔUIM proteasome | $1.47 \pm 0.03$ | $5.06 \pm 0.14$ |
| ARR-Pru-ΔUIM proteasome + 10 μM K48-Ub<sub>4</sub> | $1.14 \pm 0.02$ | $4.58 \pm 0.10$ |
| Rpn10<sup>EKK</sup> proteasome | $0.87 \pm 0.01$ | $6.61 \pm 0.20$ |
| ARR-Pru-ΔUIM-A89F proteasome | $1.40 \pm 0.03$ | $5.32 \pm 0.13$ |
| ARR-Pru-ΔUIM-A89F proteasome + 10 μM K48-Ub<sub>4</sub> | $0.54 \pm 0.01$ | $4.22 \pm 0.10$ |

**Supplementary Table 2:**
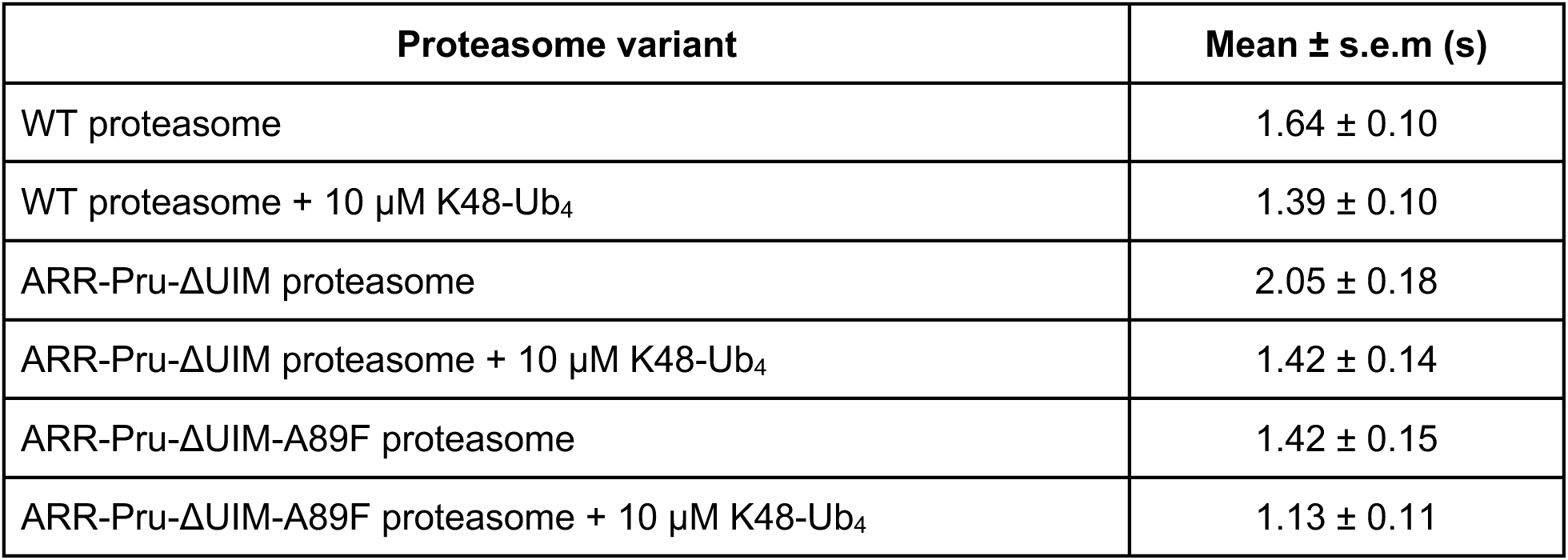
Substrate tail-insertion and engagement kinetics. The mean values of the time constants for tail insertion and engagement of SspB-delivered I27V15P substrate were determined by fitting the dwell time distribution of the tail insertion phase in the FRET-based substrate processing assay with Gamma function for wild-type proteasome and various proteasome variants in the absence and presence of unanchored K48-linked tetraubiquitin chains.

## Material and Methods

### Yeast strains

| **Name** | **Genotype** | **Source** |
| --- | --- | --- |
| yAM54 | MATa his3Δ200, leu2-3,112 lys2-801, trp1Δ63, ura3-52, PRE1-3xFLAG::KANMX | <sup>12</sup> |
| yAM80 | MATa ade2-1,15 leu2-3,112 trp1-1, ura3-1, can1-100, bar1, PRE1-Avi-HRV-3xFLAG::HIS3MX | <sup>36</sup> |

### Recombinant DNA

| **Name** | **Description** | **Source** |
| --- | --- | --- |
| pAM80 | pACYCDuet-1_Sem1-Hsp90 | <sup>12</sup> |
| pAM81 | pETDuet-1_Rpn1_Rpn2_Rpn13 | <sup>12</sup> |
| pAM83 | pACYCDuet-1_Nas2_Nas6_Hsm3_Rpt14_RILrare-tRNAs | <sup>12</sup> |
| pAM85 | pETDuet-1_Rpn9_Rpn11_Rpn8_MBP-HRV-Rpn6_Rpn5 | <sup>12</sup> |
| pAM86 | pCOLADuet-1_His6-HRV-Rpn12_Rpn7_Rpn3 | <sup>12</sup> |
| pAM87 | pUltra_AzFRS.2.t1_UAG-tRNA | <sup>12</sup> |
| pAM88 | pCOLADuet-1_FLAG-Rpt1[I191TAG]_Rpt2_His6-Rpt3_Rpt4_Rpt5_Rpt6 | <sup>12</sup> |
| pAM89 | pCOLADuet-1_FLAG-Rpt1_Rpt2_His6-Rpt3_Rpt4_Rpt5[Q49TAG]_Rpt6 | <sup>12</sup> |
| pAM210 | pCOLADuet-1_FLAG-Rpt1_sspB-Rpt2_His6-Rpt3_Rpt4_Rpt5_Rpt6 | <sup>36</sup> |
| pAM239 | pACYC-His6-Rpn10 | <sup>23</sup> |
| pAM314 | pETDuet-1_Rpn9[F2TAG]_Rpn11_Rpn8_MBP-HRV-Rpn6_Rpn5 | <sup>36</sup> |
| pAM315 | pACYC-His6-Rpn10[ΔUIM] | This paper |
| pAM321 | pETDuet-1_Rpn1[D541A, D548R, E552R]_Rpn2_Rpn13 | This paper |
| pAM322 | pETDuet-1_Rpn1_Rpn2_Rpn13[E41K, E42K, L43A, F45A, S93D] | This paper |
| pAM323 | pETDuet-1_Rpn1[D541A, D548R, E552R]_Rpn2_Rpn13[E41K, E42K, L43A, F45A, S93D] | This paper |
| pAM326 | pACYC-His6-Rpn10[R23E, D31K, E68K] | This paper |
| pAM341 | pETDuet-1_Rpn9_Rpn11[A89F]_Rpn8_MBP-HRV-Rpn6_Rpn5 | This paper |
| pAM342 | pETDuet-1_Rpn9[F2TAG]_Rpn11[A89F]_Rpn8_MBP-HRV-Rpn6_Rpn5 | This paper |
| pAM345 | pETDuet-1_Rpn9_Rpn11[A89I]_Rpn8_MBP-HRV-Rpn6_Rpn5 | This paper |
| pAM347 | pETDuet-1_Rpn9_Rpn11[L132R]_Rpn8_MBP-HRV-Rpn6_Rpn5 | This paper |
| pAM51 | pETDuet-1: Rpn8(1-179)-PreScission-Strep, H6-PreScission-Rpn11(2-239) | <sup>45</sup> |
| pAM52 | pETDuet-1: Rpn8(1-179)-PreScission-Strep, H6-PreScission-Rpn11(2-239, A89F) | <sup>45</sup> |
| pAM369 | pETDuet-1: Rpn8(1-179)-PreScission-Strep, H6-PreScission-Rpn11(2-239, A89I) | This paper |
| pAM370 | pETDuet-1: Rpn8(1-179)-PreScission-Strep, H6-PreScission-Rpn11(2-239, L132R) | This paper |
| pAM75 | pET28a MWG-Ubiquitin-G77C78 | 45 |

## Methods

### Protein Purification

Protein purification steps were done at 4⁰C unless otherwise indicated.

#### Purification of the 20S core

The 20S core particle was purified from *S. cerevisiae* as described previously ^52^. For bulk enzymatic assay, the Pre1-3xFLAG yeast strain yAM54 was used for purification. For single-molecule assay, the Pre1-Avi-HRV-3xFLAG yeast strain yAM80 was used ^36^. Yeast cultures were grown at 30⁰C in YPD media for 3 days. The cells were pelleted and resuspended in core lysis buffer (60mM HEPES pH 7.6, 400mM NaCl, 100mM KCl, 1mM EDTA), popcorned into liquid nitrogen and lysed under liquid nitrogen using a 6875 Freezer/Mill high-capacity cryogenic grinder (SPEX SamplePrep). The cryogrounded yeast powder was resuspended in core lysis buffer supplemented with 0.2% NP-40 (ThermoFisher) and thawed in the room temperature water bath. The lysate was clarified by centrifuging at 30,000 xg for 45 min. The Flag-tagged core particles were purified from the clarified lysate by incubating with M2 anti-FLAG resin (MilliporeSigma) for 1.5hr on a gentle rotator. The beads were transferred to a gravity flow column, washed with core lysis buffer and then core lysis buffer supplemented with 500mM NaCl to remove the bound 19S regulatory particle. The core particle was eluted from the resin using gel filtration buffer (GF: 30mM HEPES pH 7.6, 50mM NaCl, 50mM KCl, 10mM MgCl_2_, 5% glycerol) supplemented with 0.3mg/mL 3xFLAG peptide, concentrated and further purified by size-exclusion chromatography with a Superose 6 Increase 10/300 column (Cytiva) equilibrated in GF buffer supplemented with 0.5mM TCEP. For biotinylated 20S core particle, the beads were washed with further washed with biotinylation buffer (10mM Tris pH8.0, 25mM NaCl, 10mM MgCl_2_) before elution and eluted with biotinylation buffer supplemented with 0.3mg/mL 3x FLAG peptide. The eluted core particles were concentrated and biotinylated in biotinylation buffer supplemented with 100µM D-biotin, 1.5mM BirA biotin ligase (see below) and 10mM ATP overnight before further purification with size-exclusion chromatography.

#### Purification of the recombinant base subcomplex

The recombinant base subcomplex was purified from *E. Coli* as described previously ^12,53^. Plasmids containing yeast base subunits (Rpt1-6 including an amber codon TAG at the desired amino acid position for site-specific unnatural amino acid incorporation, Rpn1, Rpn2, and Rpn13), four base assembly chaperones (Rpn14, Hsm3, Nas2 and Nas6), and unnatural amino acid 4-azido-L-phenylalanine (AzF) tRNA synthetase/tRNA pair (for base subunits with amber codon TAG) were transformed into BL21-Star (DE3) (Invitrogen) using electroporation. Transformed bacterial cells were grown at 37⁰C in 3 liters of dYT media to an OD_600_ between 0.6 and 0.7, then pelleted, and resuspended in 1 liter of unnatural amino acid media (24 g yeast extract, 20 g tryptone, 50% glycerol buffered with 17 mM monopotassium phosphate and 72 mM dipotassium phosphate) supplemented with 2mM AzF (Acrotein ChemBio Inc). Resuspended bacterial cells were incubated at 37⁰C for 30 min before inducing with 1mM isopropyl-β-d-thiogalactopyranoside (IPTG) for 5 hours at 30°C, and overnight at 16°C. For purifying base subcomplex without unnatural amino acids, the resuspension step in unnatural amino acid media was skipped. Induced cells were pelleted by centrifugation, and resuspended in lysis buffer (60mM HEPES pH 7.6, 100mM NaCl, 100mM KCl, 10mM MgCl_2_, 5% glycerol) supplemented with 20mM imidazole, 1 mM ATP, lysozyme, benzonase (Novagen), and protease inhibitors (aprotinin, leupeptin, pepstatin, and AEBSF). The cells were lysed by sonication, the lysate was clarified by centrifuging at 30,000 xg for 30 min. The base subcomplex was purified using double affinity tags on Rpt subunits. First, the His-Rpt3 containing complexes were purified by batch binding to Ni agarose beads (ThermoFisher) in lysis buffer supplemented with 20 mM imidazole, 1 mM ATP for 45 min, and eluting with lysis buffer supplemented with 1 mM ATP and 250 mM imidazole. Then the fully assembled base subcomplexes containing Flag-Rpt1 were purified by column binding to M2 anti-FLAG resin (MilliporeSigma), and eluting with GF buffer supplemented with 0.3mg/mL Flag peptide, MDYKSHDGDYKDHDIDYKDDDDKG (Genscript Bio Corp). The eluted base complexes were concentrated, incubated with 150 μM 5,5′-dithiobis-(2-nitrobenzoic acid) (DTNB) for 10 min at room temperature to reversibly block surface-exposed cysteines before the AzF (incorporated in place of Q49 of Rpt5 or I191 of Rpt1) was reacted with 300 μM dibenzocyclooctyne (DBCO)– conjugated LD655 dye (a sulfo-Cy5 derivative from Lumidyne Technologies; the DBCO modification is custom synthesis) at 4°C overnight. The reaction was quenched with 1 mM free AzF, followed by addition of 5 mM dithiothreitol (DTT) to reverse the DTNB modification of cysteines. The labeled base subcomplexes were further purified by size-exclusion chromatography with a Superose 6 Increase 10/300 column (Cytiva) equilibrated in GF buffer supplemented with 0.5mM ATP and 0.5mM TCEP. Base concentrations were determined by Bradford assay, and labeling efficiencies were determined to be ∼70 to 90% according to absorbance of the fluorophore measured using Nanodrop.

#### Purification of the recombinant lid subcomplex

The recombinant lid subcomplex was purified from *E. Coli* as described previously ^12,53^. Plasmids containing yeast lid subunits (Rpn3, Rpn5, Rpn6, Rpn7, Rpn8, Rpn9, Rpn11, Rpn11, and Sem1), and unnatural amino acid 4-azido-L-pheylalanine (AzF) tRNA synthetase/tRNA pair (for base subunits with amber codon TAG) were transformed into BL21-Star (DE3) (Invitrogen) using electroporation. Transformed bacterial cells were grown at 37⁰C in 3 liters of dYT media to an OD_600_ between 0.6 and 0.8, then pelleted, and resuspended in 1 liter of unnatural amino acid media (24g yeast extract, 20g tryptone, 50% glycerol buffered with 17mM monopotassium phosphate and 72mM dipotassium phosphate) supplemented with 2mM AzF (Acrotein ChemBio Inc). Resuspended bacterial cells were incubated at 37⁰C for 30 min before inducing with 1mM isopropyl-β-d-thiogalactopyranoside (IPTG) for 5 hours at 30°C, and overnight at 16°C. For purifying lid subcomplex without unnatural amino acids, the resuspension step in unnatural amino acid media was skipped. Induced cells were pelleted by centrifugation, and resuspended in lysis buffer supplemented with 20 mM imidazole, lysozyme, benzonase (Novagen), and protease inhibitors (aprotinin, leupeptin, pepstatin, and AEBSF). The cells were lysed by sonication, the lysate was clarified by centrifuging at 30,000 xg for 30 min. The lid subcomplex was purified using double affinity tags on lid subunits. First, the His-Rpn12 containing complexes were purified by batch binding to Ni agarose beads (ThermoFisher) in lysis buffer supplemented with 20 mM imidazole for 45 min, and eluting with lysis buffer supplemented with 250 mM imidazole. Then the fully assembled base subcomplexes containing maltose-binding protein (MBP) fused Rpn6 were purified by column binding to amylose resin (NEB), and eluting with GF buffer supplemented with 10mM maltose. The eluted lid complexes were concentrated, incubated with HRV-protease for 1 hour at room temperature to cleave the MBP tag, then with 150 μM 5,5′-dithiobis-(2-nitrobenzoic acid) (DTNB) for 10 min at room temperature to reversibly block surface-exposed cysteines before the AzF (incorporated in place of Q49 of Rpt5 or I191 of Rpt1) was reacted with 300 μM dibenzocyclooctyne (DBCO)–conjugated LD555 dye (a sulfo-Cy3 derivative from Lumidyne Technologies; the DBCO modification is custom synthesis) at 4°C overnight. The reaction was quenched with 1 mM free AzF, followed by addition of 5 mM dithiothreitol (DTT) to reverse the DTNB modification of cysteines. The labeled lid subcomplexes were further purified by size-exclusion chromatography with a Superose 6 Increase 10/300 column (Cytiva) equilibrated in GF buffer supplemented with 0.5 mM TCEP. Lid concentrations were determined using Nanodrop, and labeling efficiencies were determined to be ∼70 to 90% according to absorbance of the fluorophore measured using Nanodrop.

#### Purification of the recombinant Rpn10 subunit

The recombinant Rpn10 subunit was purified from *E. Coli* as described previously ^36^. Plasmid containing yeast Rpn10 subunit with N-terminal His tag was transformed into BL21-Star (DE3) (Invitrogen) using heat shock. Transformed bacterial cells were grown at 37⁰C in 2 liters of dYT media to an OD_600_ between 0.6 and 0.8, then induced with 0.5 mM isopropyl-β-d-thiogalactopyranoside (IPTG) for 4 hours at 37°C. Induced cells were pelleted by centrifugation, and resuspended in lysis buffer supplemented with 20mM imidazole, lysozyme, benzonase (Novagen), and protease inhibitors (aprotinin, leupeptin, pepstatin, and AEBSF). The cells were lysed by sonication, the lysate was clarified by centrifuging at 30,000 xg for 30 min. His-tagged Rpn10 subunits were purified from the clarified lysate by batch binding to Ni agarose beads in lysis buffer supplemented with 20 mM imidazole for 45 min, and eluting with lysis buffer supplemented with 250 mM imidazole. Eluted Rpn10 subunits were concentrated, and further purified by size-exclusion chromatography with a Superdex 75 16/60 column (Cytiva) equilibrated in GF buffer supplemented with 1 mM DTT. Lid concentrations were determined using Nanodrop.

#### Purification of substrates

The titin I27 substrate was purified from *E. Coli* as described previously ^36^. Plasmid containing titin I27 substrate with N-terminal Glycine-Glycine-Glycine (GGG) tag for fluorescent labeling and C-terminal chitin-binding domain (CBD) was transformed into BL21-Star (DE3) (Invitrogen) using heat shock. Transformed bacterial cells were grown at 30⁰C in 2 liters of M9 minimal media to an OD_600_ 0.6, then induced with 0.5 mM isopropyl-β-d-thiogalactopyranoside (IPTG) for 3 hours at 30 °C. Induced cells were pelleted by centrifugation, and resuspended in titin lysis buffer (60mM HEPES pH 7.6, 100mM NaCl, 1 mM EDTA, 5% glycerol) supplemented with benzonase (Novagen), and protease inhibitors (aprotinin, leupeptin, pepstatin, and AEBSF). The cells were lysed by sonication, the lysate was clarified by centrifuging at 30,000 xg for 30 min. The clarified lysate was incubated with chitin resin (NEB) in the lysis buffer for 1 hour. Substrate bound resin was washed with lysis buffer supplemented with 550 mM NaCl and 0.1% Triton X-100, and incubated with cleavage buffer (30mM HEPES pH 8.5, 100mM NaCl, 1mM EDTA, 5% glycerol, 50mM DTT) overnight to cleave titin substrate from chitin resin. The flowthrough containing the cleaved substrates was collected, flowed over fresh chitin resin to remove any uncleaved protein, concentrated and further purified by size-exclusion chromatography with a Superdex 75 16/60 column (Cytiva) equilibrated in GF buffer supplemented with 0.5 mM TCEP. For bulk degradation assay, substrates were covalently labeled with the fluorescein (FAM) peptide FAM-HHHHHHLPETGG (Genscript) via sortase reaction to its N-terminal GGG tag. Briefly, 100 μM purified substrates were incubated with 20 μM sortase and 500 μM FAM peptide in GF buffer supplemented with 10 mM CaCl_2_ and 1 mM DTT for 2 hours at room temperature. Labeled substrates were enriched using Ni-NTA agarose to ensure 100% labeling efficiency, and further purified by size-exclusion chromatography with a Superdex 75 10/300 column (Cytiva) equilibrated in GF buffer supplemented with 0.5 mM TCEP. For single molecule assay, substrates were labeled with LD555-maleimide dye (a sulfo-Cy3 derivative from Lumidyne Technologies) at the engineered cysteine position in the C-terminal flexible initiation region. 100 μM purified substrates were buffer exchanged into maleimide labeling buffer (30mM HEPES pH 7.2, 150mM NaCl, 1 mM EDTA), and then labeled with 100 μM LD555-maleimide dye at room temperature for 1 hour. The labeling reaction was quenched by incubating with 10 mM DTT for 15 minutes at room temperature. Labeled substrates were further purified by size-exclusion chromatography with a Superdex 75 10/300 column (Cytiva) equilibrated in GF buffer supplemented with 0.5 mM TCEP.

#### Ubiquitination of substrates

Substrates were poly-ubiquitinated as described previously (Bard Cell 2019). Briefly, 10 μM substrate, 1 μM mE1, 2.5 μM UbcH7, 2.5 μM Rsp5, 1 mM Ubiquitin, and 10 mM ATP in GF buffer were reacted for 1 hour at 30 °C. Ubiquitination was assessed by a 4-20% SDS-PAGE (Tris-Glycine, Biorad). For mono-ubiquitinated substrate, wild-type ubiquitin was replaced with ubiquitin K63R ubiquitin to prevent chain formation. Mono-ubiquitinated substrate was separated from un-ubiquitinated substrate using a Superdex S75 increase 10/300 (Cytiva), concentrated, and the concentration was determined using the extinction coefficient of the fluorescein dye.

#### Purification and synthesis of K48-linked tetraubiquitin chains

K48 tetraubiquitin chains were synthesized and purified as described previously (Dong, Structure 2011). 1 mM WT Ubiquitin, 1 μM mE1, 2.5 μM Cdc34, and 10 mM ATP were reacted in GF buffer overnight at 37 degrees C. Tetra-ubiquitin was separated from the other lengths on a Resource S cation exchange (Cytiva) using a 0 - 1 M NaCl gradient in 50 mM sodium acetate buffer pH 4.5. Tetraubiquitin was further purified on a Superdex S75 increase 10/300 column in GF buffer (Cytiva).

#### Mutation, expression, and purification of Rpn11/Rpn8

Mutations to Rpn11 were introduced using around-the-horn PCR. The Rpn11/Rpn8 heterodimer was expressed and purified as previously described ^24^. Rpn11/8 was expressed in BL21 (DE3) cells grown at 37 degrees C in Terrific Broth and induced with 1 mM IPTG overnight at 18 degrees C. These cells were sonicated and then clarified by 15,000xg centrifugation and purified over a HiTrap Ni NTA column followed by an Superdex 200 16/60 size exclusion column (Cytiva) in GF buffer. Protein concentrations were determined by the absorbance at 280 nm.

#### Rpn11/Rpn8 deubiquitination assays

Deubiquitination assays were performed as described ^54^. 0.5 μM of Rpn11/8 was incubated with 100 μM Ubiquitin-TAMRA in a 384-well low-volume black flat bottom plate (Corning, #3820) and the fluorescence polarization of TAMRA was monitored using a CLARIOstar Plus plate reader (BMG Labtech) at 30°C.

#### Purification of biotin ligase

The recombinant BirA biotin ligase was purified from *E. coli*. BL21-Star (DE3) (Invitrogen) cells were transformed with the plasmid containing *E. coli* BirA with N-terminal His and MBP tags using heat shock. Transformed cells were grown at 37⁰C in 2 liters of dYT media to an OD_600_ between 0.6 and 0.8, then induced with 0.5 mM isopropyl-β-d-thiogalactopyranoside (IPTG) overnight at 18°C. Induced cells were pelleted by centrifugation, and resuspended in lysis buffer supplemented with 20mM imidazole, lysozyme, benzonase (Novagen), and protease inhibitors (aprotinin, leupeptin, pepstatin, and AEBSF). The cells were lysed by sonication, the lysate was clarified by centrifuging at 30,000 xg for 30 min. His-and MBP-tagged BirA was purified from the clarified lysate by batch binding to Ni agarose beads in lysis buffer supplemented with 20 mM imidazole for 45 min, eluting with lysis buffer supplemented with 250 mM imidazole, binding to the amylose resin (NEB), and eluting with GF buffer supplemented with 10mM maltose. Eluted BirA were concentrated, and further purified by size-exclusion chromatography with a Superdex 200 16/60 column (Cytiva) equilibrated in GF buffer supplemented with 1 mM DTT.

#### Fluorescence polarization-based multiple-turnover degradation assay

Reconstituted proteasome mix was prepared at 2x concentration by mixing and incubating 100 nM core particle, 800 nM SspB-fused base, 1.2 μM lid, and 2 μM Rpn10 at room temperature for 10 min in GF buffer supplemented with an ATP regeneration system (0.03 mg/mL creatine kinase and 16 mM creatine phosphate). Substrate mix was prepared at 2x concentration (20 μM) in GF buffer supplemented with 1 mg/mL bovine serum albumin and 10 mM ATP. For experiments with unanchored K48-linked tetraubiquitin chains, 20 μM chains were added in the substrate mix. The multiple-turnover degradation reaction was initiated by mixing 7 μl of 2x reconstituted SspB-fused proteasome and 7 μl of 2x substrate mix. Ten microliters of the reaction was immediately transferred to a 384-well low-volume black flat bottom plate (Corning, #3820) prewarmed to 30°C, and degradation was monitored by the loss of fluorescence polarization signal from fluorescein (FAM)–labeled substrates in a CLARIOstar Plus plate reader (BMG Labtech) at 30°C. Rates for multiple-turnover degradation were determined by linear regression of the initial change in polarization and normalizing the fit with the measured differences in fluorescence polarization signals of undegraded substrates and substrates fully degraded by 0.1 μg/μl chymotrypsin.

#### TIRF microscopy

Single-molecule imaging was performed with an inverted Eclipse Ti2 microscope (Nikon) equipped with a TIRF 60x 1.49 N.A. oil immersion objective (Nikon) and a LUNF laser launch with 488 nm, 532 nm, 561 nm and 640 nm laser lines (Nikon). The emitted signals were splitted into two halves using a W-view Gemini imaging splitting optics (Hamamatsu) and detected on an electron multiplying CCD camera (Andor Technology, iXon Ultra 897). Spatial registration of the split emissions channels was performed by overlaying the imaging signals from 0.1 µm Tetraspeck microspheres (Thermofisher). Illumination and image acquisition is controlled by NIS Elements Advanced Research software (Nikon).

#### Single-molecule conformational dynamics and substrate-processing assays

Single-molecule conformational dynamics and substrate processing assays were performed as described previously ^36^. The imaging chamber was assembled by sticking low-density PEG-biotin-coated coverslips (Microsurfaces) onto Superfrost microscope slides (Fisherbrand) using double-sided tapes (Scotch). 0.05 mg/mL NeutrAvidin (Thermofisher) in the GF buffer was flowed into the imaging chamber and incubated for 3 minutes. Excess NeutrAvidin was washed twice with the assay buffer (GF buffer supplemented with 0.4 mg/mL BSA, 1.2 mM Trolox, 0.4 mM 2-mercaptoethanol, 2 mM ATP, 0.015 mg/mL creatine kinase and 8 mM creatine phosphate). Reconstituted proteasome mix was prepared by mixing and incubating 500 nM core particle, 400 nM recombinant base labeled with LD655 on the Rpt5 subunit for conformational dynamics assay or on the Rpt1 subunit for substrate processing assay, 600 nM recombinant lid labeled with LD555 on the Rpn9 subunit for conformational dynamics assay, and 1 μM Rpn10 at room temperature for 10 min in GF buffer supplemented with an ATP regeneration system [0.03 mg/mL creatine kinase and 16 mM creatine phosphate]. Reconstituted proteasome was diluted to 1 nM to 100 pM, flowed into the imaging chamber and incubated for 5 minutes. The unbound proteasome subunits were washed away twice with the assay buffer supplemented with 300 nM Rpn10. The final imaging buffer containing the assay buffer, 300 nM Rpn 10, and oxygen scavenging system (5 mM protocatechuic acid and 100 nM protocatechuate-3,4-dioxygenase) was flowed in before imaging. For experiment with unanchored K48-linked tetraubiquitin chains, 100 µM chain was supplemented in the final imaging buffer. For substrate processing assays, 50 nM of LD555- labeled titin substrate was added to the final imaging buffer. The imaging was performed by first taking a single-frame snapshot of the acceptor (LD655) signal by exciting with 50 ms exposure of the 640 nm laser using the TIRF microscopy setup described above. The donor (LD555) fluorophores were exposed for 50 ms with the 532 nm laser, and imaged at a frame rate of 51.7 ms for 2000 frames.

Single-molecule conformational dynamics and substrate processing data analysis were performed as described previously ^36^. Briefly, single-molecule conformational dynamics data was analyzed using the Spartan software package ^55^. The donor fluorescence, acceptor fluorescence and apparent FRET efficiency traces were extracted using the gettraces function of the Spartan software package. The apparent FRET efficiency traces of singly-capped and singly-labeled proteasomes were selected for further analysis by setting the total fluorescence intensity threshold in the *autotrace* function and selecting a single photobleaching step by manual inspection. These traces were analyzed using a two-state hidden Markov model in the *ebFRET*. The dwell times of s1 state and non-s1 state were extracted and fit to a single exponential function in Prism (GraphPad). Single-molecule processing data was analyzed using a custom-built MATLAB TIRFexplorer app ^36^. The maximum intensity projection of the acceptor channel was first generated and the coordinates of the acceptor spots, which indicates the FRET events, were recorded in ImageJ. The donor fluorescence, acceptor fluorescence and apparent FRET efficiency traces were extracted using the TIRF explorer app by feeding the coordinates of the acceptor spots. These traces were inspected manually and scored for unsuccessful substrate binding and successful substrate processing events as described previously ^36^. The event where an increase in donor fluorescence colocalized with acceptor fluorescence followed by a single-step photobleaching step for both donor and acceptor signals was scored as unsuccessful binding events. The event where an increase in donor fluorescence colocalized with acceptor fluorescence resulting in a medium FRET value followed by an increase to a high-FRET phase, a donor-fluorescence recovery and extended donor-fluorescence dwell was scored as successful substrate processing events. Capture success was calculated as the percentage of successful processing events relative to the total number of substrate binding (both successful and unsuccessful). The tail insertion times of successful substrate capture events were further extracted by recording the time from the appearance of a donor signal to the point of the first peak of high FRET efficiency.

#### Quantification and statistical analyses

Details about the statistical analyses for individual experiments, including the statistical tests and exact values of N (number of technical replicates), can be found in the figure legends. Error bars represent the mean ± S.D.

